# EEG Microstate Dynamics During Immersive VR Earthquake Exposure: Signatures of Threat Reactivity, Recovery, and Resilience

**DOI:** 10.64898/2026.07.28.741178

**Authors:** Marius A. Dragu, Ciprian Fǎcǎeru, Magdalena Țurcanu, Cosmina A. Duțicǎ, Alexandru Bîcu, Olguța Barizi, Octavian F. Mirica, Alexandra Sofonea, Gabriela Niculescu, Miralena I. Tomescu

## Abstract

Adapting to natural disasters requires rapid and flexible regulation of emotional and physiological responses, yet the large-scale brain dynamics supporting this process remain poorly understood. Using an immersive virtual reality (VR) earthquake simulation, we investigated real-time neural adaptation to acute environmental threat and its modulation by resilience.

Seventy-six participants were randomly assigned to a VR earthquake condition (VR-ES, n = 38) or a control VR condition (VR-CTRL, n = 38). Resting-state EEG was recorded before and after the VR experience, and continuous mobile EEG was recorded during the simulation. EEG microstate analysis was used to characterize large-scale brain dynamics associated with threat exposure and recovery.

Exposure to the simulated disaster induced distinct alterations in microstate dynamics, with opposing changes in microstates C (p<0.001, r_b_=0.55) and D (p<0.001, r_b_=0.52) observed specifically in the VR-ES group. Moreover, our results confirm that distinct neural patterns are activated during the recovery resting-state phase following VR-ES. Critically, resilience and childhood adversity moderated the patterns of associations with emotional recovery, such that individuals with higher resilience exhibited more adaptive microstate configurations associated with improved recovery outcomes.

These findings suggest that flexible reorganization of large-scale brain states supports adaptation to acute environmental stress and that resilience shapes the neural dynamics underlying recovery. This work highlights the value of ecologically valid VR paradigms combined with mobile neurophysiology for understanding individual differences in responses to environmental threat and their adaptive neural mechanisms.

## Introduction

Natural disasters vividly reveal the volatility of our environment and the limits of human adaptability, underscoring the critical importance of elucidating the neural mechanisms that govern rapid reactivity and recovery in the face of extreme threat (Knez et al., 2021; Mortimer et al., 2026; Vardy & Atkinson, 2019; Waters et al., 2022; Zhao et al., 2024). These challenges are being amplified by climate change, a major driver of the increasing frequency and severity of natural disasters (Banholzer et al., 2014; Dowdy et al., 2019). Between 1998 and 2017, such events claimed the lives of approximately 1.3 million people worldwide and affected an estimated 4.4 billion individuals through injury, displacement, homelessness, or the need for emergency assistance (Wallemacq et al., 2018).

Despite their profound impact, how humans prepare, rapidly adapt during natural disasters, and recover immediately after remains understudied. This is largely due to the ethical, practical, and methodological challenges inherent in studying neural and physiological responses during real-world extreme events.

To overcome this gap in the literature, immersive virtual reality (VR) environments, particularly CAVE-based virtual reality, offer a powerful solution by enabling controlled, safe simulation of natural disasters. VR represents an ecologically valid and controllable method for investigating cognitive and affective processes under dynamic, interactive, and realistic conditions (Nuojua et al., 2024; Parsons, 2015; Schöne et al., 2023; Zanier et al., 2018). Unlike traditional laboratory tasks, which often rely on simplified stimuli and constrained motor responses, VR enables the presentation of multimodal and immersive scenarios that closely replicate real-life situations, thereby enhancing the ecological validity of the findings (Ascone et al., 2025; Deng et al., 2024; Hameed et al., 2023; Hanshans et al., 2024; Huijsmans et al., 2025; Mioni & Pazzaglia, 2023; Oleksy et al., 2024). CAVE-VR systems facilitate the design of complex multimodal stimuli while reducing external distraction and broaden the behavioral repertoire available for investigation, enabling the observation of whole-body movements, gaze patterns, and reflexive actions in response to emotional stimuli or stressors (Ma et al., 2025; Schöne et al., 2023). More importantly, the CAVE-VR systems can be easily integrated with electrophysiology, including mobile brain EEG studies (Gramann, 2024). Recent studies have demonstrated that EEG research conducted in immersive environments, such as virtual reality, augmented reality, and mixed reality, offers valuable insights into fundamental cognitive functions, including attention, cognitive load, spatial awareness, and brain-body interaction (Akounach et al., 2026; Denzer et al., 2024; Kawai et al., 2024; Ma et al., 2025). The flexibility of VR scenarios allows precise manipulation of environmental parameters, which can be synchronized with EEG recordings to assess brain activity in real time (Li et al., 2024).

EEG microstates (MSs) enabling real-time characterization of the reorganization of large-scale neural networks with fast temporal dynamics, can capture intra- and inter-individual variability in cognitive and socio-emotional adaptation to imagined, real life, and VR environments (Denzer et al., 2024; Deolindo et al., 2021; Nazare & Tomescu, 2024; Tomescu et al., 2022). EEG MSs capture the coordinated activity of distributed neural generators and provide a temporally precise window into network-level processing steps of information across a variety of brain states (Michel & Koenig, 2018; Tarailis et al., 2024).

Moreover, accumulating evidence indicates that EEG microstates represent electrical signatures of resting-state brain networks (Britz et al., 2010; Custo et al., 2017; Musso et al., 2010; Yuan et al., 2012). However, definitive one-to-one mappings between microstates and their neural generators remain unresolved, highlighting the need for further research. Generally, microstates A and B are associated with the visual and auditory/ language sensory cortices and with arousal-related processes. Microstate B is particularly involved in visual processing, including self-referential functions and autobiographical memory, and its role extends beyond visual stimulation through interactions with other microstates (Antonova et al., 2022; Michel & Koenig, 2018; Milz et al., 2016; Seitzman et al., 2017; Tarailis et al., 2024). Following acute stress induction paradigms like the Trier Social Stress Test (TSST), coverage and duration of MS B decreased, while other studies reported increased presence also during positive affective states, suggesting MS B might be involved in the rapid recovery of affective state with important clinical implications (Chivu et al., 2024; Hu et al., 2021; Nazare & Tomescu, 2024). Microstates C, D, and E have been linked to key regions of top-down functional networks, such as the default mode network (DMN), the dorsal attention network (DAN), and the salience network (SN) (Michel & Koenig, 2018). Microstate C is involved in the processing of personally meaningful information; microstate D is closely related to higher-order cognitive functions such as working memory and attention; and microstate E is associated with interoceptive and emotional processing, including the detection of salience and emotional relevance (Antonova et al., 2022; Deolindo et al., 2021; Faber et al., 2017; Michel & Koenig, 2018; Schiller et al., 2019; Tarailis et al., 2024). During emotional regulation and recovery of affective states, increased coverage and occurrence of MS C were observed following acute stress (Hu et al., 2021), and a decreased MS C duration of self-generated negative emotional regulation was observed (Nazare & Tomescu, 2024). In contrast, a reversed pattern of MS D increased presence was observed during self-generated negative emotional states (Nazare & Tomescu, 2024). Importantly, these stress-induced changes were linked to physiological responses: cortisol levels measured 10 minutes after the TSST were significantly elevated and negatively associated with the transition probability between MS C and D (Hu et al., 2021), further strengthening the role of higher-order top-down MS in affective state regulation and stress recovery. MS E, among the new ones to be investigated in the MS literature, also shows an association with affective state regulation with reduced presence during negative emotional states (Nazare & Tomescu, 2024). Most importantly, MS E, also found in literature as MS C, given the high spatial correlation with MS C topography, has also been associated with immersive states of subjective experience of reality, bizarreness in dream-like VR experiences (Denzer et al., 2024). Finally, MSs C, D, and E have also been shown to modulate complex real-life tasks like flying a helicopter, with MS C and D increased and MS E decreased activity compared to pre- and post-resting states (Deolindo et al., 2021).

The primary objective of this study is to examine how an ecologically valid VR simulation of a natural disaster modulates neurohormonal and affective reactivity, and brain network dynamics via the temporal dynamics of EEG microstate structures by using an integrative approach combining mobile brain EEG within CAVE-VR, providing novel insights into how humans adapt in real time to ecologically valid, immersive simulations of natural disasters. Examining spatiotemporal EEG microstate dynamics and their associations with affective regulation and hypothalamic–pituitary–adrenal (HPA) axis via cortisol, during and following an immersive session simulating an earthquake (VR-ES), we aim to determine intra- and inter-individual variability in VR-ES response as a function of resilience traits and prior adverse childhood experiences. To the best of our knowledge, this study represents the first multimodal investigation to link EEG microstate dynamics with affective and endocrine responses in the context of a simulated natural disaster, thereby identifying neurophysiological markers of resilience to adverse natural disaster experiences.

## Methods

### Ethical considerations

All methods and experiments received approval from the Ethics Committee of the National University for Theatre and Film I.L Caragiale, Bucharest and adhered to the Declaration of Helsinki guidelines. All participants gave written informed consent to participate.

### Experimental design

This study investigates how the VR experience modulates the resting-state EEG, cortisol data, and behavioural states. Participants were randomly assigned to either the VR- earthquake simulation (VR-ES) group or the VR-control (VR-CTRL) group. To control for possible effects of cortisol variation as a function of the circadian rhythm, the experiment was conducted for each participant during the same time interval, namely between 3 p.m and 6 p.m.

The VR simulation was implemented with a CAVE-VR system, which projects stereoscopic 3D images onto three walls and the floor using WUXGA (1920×1200) laser projectors (**Figure 1 a-e**). Participants wore active stereo LCD shutter glasses synchronised with the projectors. User position and orientation were tracked in real time by an integrated ART motion-tracking system, while the virtual environment was rendered by real-time simulation software, built using Unity Engine and running on a high-performance workstation. Additionally, the simulation space contains physical props: a table, a chair, a fire extinguisher, and a smartphone, co-located with the virtual scene, resulting in a mixed-reality environment that combines virtual and physical elements. The broader experimental technology stack also included a 4.1 surround system in a four-corner loudspeaker configuration with an additional subwoofer and, most importantly, concurrent physiological acquisition hardware (portable high-density EEG) synchronised with the CAVE system virtual reality application.

**Figure 1.**
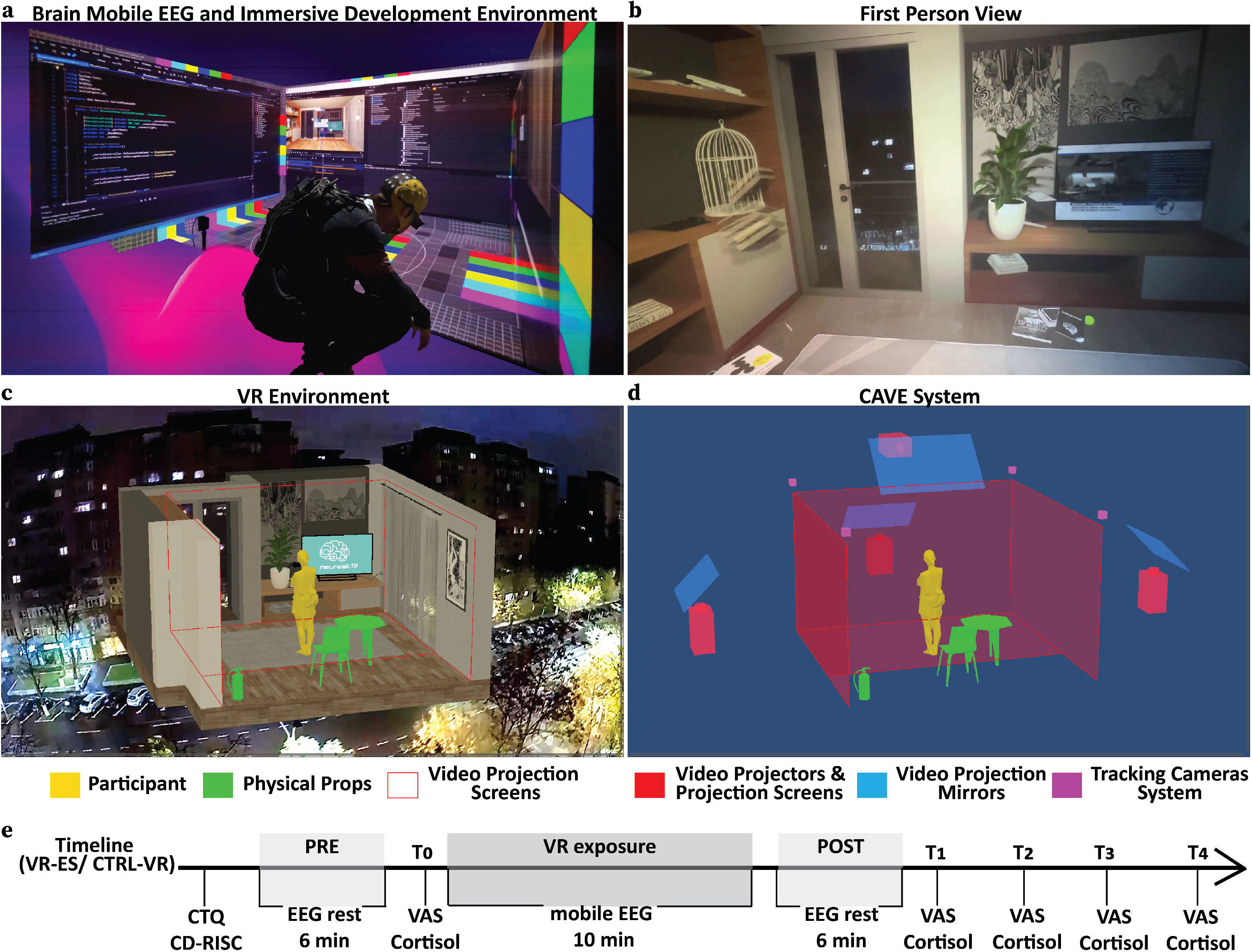
Overview of the CAVE-Based VR Environment and Experimental design. **(a)** Brain Mobile EEG and Immersive Development Environment. **(b)** First Person View: what the participant sees. **(c)** VR Environment: a high-fidelity digital model of a modern residential apartment. **(d)** CAVE-Based Immersive VR System: principal components include Video Projectors and Projection Screens, Video Projection Mirrors, and Tracking Cameras System. **(e)** Experimental design: Timeline of VR exposure (VR-ES and CTRL-ES) and data collection, including resilience traits (CD-RISC questionnaire), trauma-related traits (CTQ questionnaire), mobile EEG, behavioural states (emotion and anxiety assessed using VAS) and hormonal states (salivary cortisol). T_0_ represents the time point immediately before VR exposure; T_1_, 15 minutes after VR exposure; T_2_, 30 minutes after VR exposure; T_3_, 45 minutes after VR exposure; and T_4_, 60 minutes after VR exposure.

Participants in the VR Experiment group underwent a 10-minute immersive session in a CAVE-based virtual reality system presenting an earthquake scenario in a high-fidelity digital model of a short-term rental (Airbnb-style) apartment. After a brief exploration phase, an earthquake event began without a warning. The immersion is enhanced the simulation integrated multimodal cues: the projected imagery of the virtual environment depicted building sway and vibration; spatialized multi-channel (surround) audio reproduced structural creaks and impact sounds; animated furniture and decorative objects toppled and fell; the in-scene television displayed breaking-news coverage related to the earthquake; and a physical mobile phone carried by the participant is programmed to vibrate intermittently while playing the RO-Alert emergency tone. These elements were intended to increase perceived presence and induce an acute stress response during the unfolding event.

### Participants and dataset collection

Both participant recruitment and EEG data acquisition, as well as the design of the immersive VR CAVE experience, were carried out at the International Center for Research and Education in Innovative Creative Technologies (CINETic), the Research and Development Department of the National University of Theatre and Film “Ion Luca Caragiale” in Bucharest. A total of 76 healthy participants were included in the study, comprising 37 women and 39 men, with a mean age of 22.45 ± 5.73 years (range: 14–41 years). The descriptive demographics for the groups are: VR-ES group (N = 38; 18 women, 20 men; mean age = 23.31 ± 6.88 years) and the VR-CTRL group (N = 38; 19 women, 19 men; mean age = 21.59 ± 4.33 years). Table 1 summarises the demographic statistics for the given group.

**Table 1.** Demographic data for the analysed dataset.

| Subject | Entire group | VR-ES | VR-CTRL |
| --- | --- | --- | --- |
| N/ percent | 76 | 38/ 50% | 38/ 50% |
| Age (range) | 14-41 | 14-41 | 16-38 |
| Age (*mean* ±SD) | 22.45±5.73 | 23.31±6.88 | 21.59±4.33 |
| Independent t-Test (Mann-Whitney): $t=459$, $p=0.47$ | | | |
| Gender (*F/M*) | 37/39 | 18/20 | 19/19 |
| Chi-square test: $\chi^2=0.05$, $p=0.82$ | | | |

EEG data were acquired using the portable *eego™ sports* system (ANT Neuro), optimised for movement-related recordings. A 64-channel *waveguard original* cap, configured according to the extended international 10–20 system with online sampling at 1 kHz and CPz as the reference electrode, was employed. We collected high-density EEG signals before the VR experience (PRE-VR session, duration: 6 min), during the VR experience (duration: 10 min) and after the VR experience (POST-VR session, duration: 6 min).

Self-reported data were gathered in the same room as the PRE/ POST EEG data and included various pen-and-paper tests. We used two visual analog scales (VAS) before (T_0_) and after the VR experience at four time points (T_1_ -15 minutes after VR exposure, T_2_-30 minutes after VR exposure, T_3_-45 minutes after VR exposure, T_4_-60 minutes after VR exposure- **Figure 1e**) to assess emotion and anxiety. The VAS tests used a 10-cm unmarked scale ranging from 0 (“no negative emotions” /“no anxiety”) up to 100 (“anxiety at the highest level” / “intense negative emotions (stress)”).

The Connor-Davidson Resilience Scale (CD-RISC) is a 25-item self-report questionnaire that quantifies resilience, defined as stress-coping ability (Connor & Davidson, 2003). The responses are rated on a 5-point Likert scale from 0 (“Completely Disagree”) to 4 (“Completely Agree”). The scores for each item were then summed up and reported; the total score ranges from 0 to 100, with higher scores indicating greater resilience.

The Childhood Trauma Questionnaire (CTQ) comprises 25 items to assess trauma, a measure of traumatic childhood exposures (Bernstein et al., 2003). The items are organised on five subscales: physical abuse, emotional abuse, sexual abuse, physical neglect, and emotional neglect. Each item uses a 5-point Likert scale, ranging from 1 (“Never True”) to 5 (“Often True”), based on the occurrence of each event. Total scores vary from 25 to 125, with higher scores reflecting higher levels of childhood trauma experiences. Participants completed the CD-RISC and CTQ questionnaires at the beginning, before the PRE-EEG block recording (**Figure 1e**).

Cortisol levels were measured in saliva using Sarstedt Salivette collection devices at five time points throughout the experiment (T_0_, T_1_, T_2_, T_3_, T_4_- **Figure 1e**). Participants used a mouth swab to collect sufficient saliva. The saliva samples for cortisol analysis were stored at −20 °C until processing. Cortisol concentrations (μg/dL) were quantitatively measured using an enzyme-linked immunosorbent assay. Given that salivary cortisol exhibits a pronounced circadian rhythm, the experiment was conducted between 3 p.m and 6 p.m. for each participant.

### EEG Data Processing

In the preprocessing stage, we designed and applied a Notch filter to remove power-line interference from the raw data, followed by a 1-40 Hz Butterworth band-pass filter to retain broadband neural activity. Additionally, we manually identified and removed epochs containing movement artefacts using *Cartool* and *MNE-Python 3.9* (Brunet et al., 2011; Gramfort et al., 2013). Subsequently, in *EEGLAB* (MATLAB) (Nagabhushan Kalburgi et al., 2024; Poulsen et al., 2018), we performed Adaptive Mixture Independent Component Analysis (AMICA) to decompose the EEG signal into independent components, thereby enabling the identification and removal of noise-related sources such as eye movements or muscle activity (Artoni & Michel, 2025; Klug et al., 2024). Independent components were classified using ICLabel (Pion-Tonachini et al., 2019), which estimates the probability (*p*ICLabel) that each independent component belongs to one of seven classes: Brain, Muscle, Eye, Heart, Line Noise, Channel Noise, or Other. Non-neuronal sources were automatically removed if *p*_ICLabel_(Class {Muscle, Eye, Heart, Line Noise})>0.5, while components classified as *Other* were rejected if *p*_ICLabel_(Class=Brain) <0.2. Bad channels were interpolated using a 3D Spherical Spline method in *Cartool* (Brunet et al., 2011), and the data were subsequently re-referenced to the standard mean average. Finally, the dataset was down-sampled to 125 Hz to reduce data volume and computational complexity.

### Microstate Analysis

EEG microstate analysis involves segmenting the signal into millisecond-scale periods during which the brain activity topography remains stable. We performed EEG microstate analysis to identify the dominant maps corresponding to the different microstates and to calculate their temporal dynamics descriptors: mean duration and occurrence. Specifically, these temporal electrophysiological markers, extracted via microstate analysis, highlight VR-related changes in neural dynamics.

First, we computed the Global Field Power (GFP) (Lehmann & Skrandies, 1980) , defined as the standard deviation of the biopotentials across all electrodes in an average reference map (Mishra et al., 2020). The local maxima of GFP exhibit optimal signal-to-noise ratios in EEG signals and serve as ideal markers for identifying quasi-stable topographic patterns (Michel & Koenig, 2018; Murray et al., 2008).

To identify the most representative topographies, the K-means algorithm was applied in two steps: first at the individual level, then at the group level. Individual (subject)-level clustering was performed using EEG signals extracted from the time frames corresponding to GFP peaks as inputs. To reduce computational time while obtaining a representative set that captures sufficient variability in the data and provides a robust basis for group-level clustering, we selected the same number of individual topographies per subject-seven. This choice was guided by the maximum number of distinct microstate topographies (A–G) consistently referenced in the literature (Tarailis et al., 2024). To automate the individual clustering process and optimise runtime, based on the MNE Python library (Gramfort et al., 2013), we developed a pipeline that extracts the most representative topographies from each recording as a 7 (dominant topographies) x 64 (electrodes) array, associated with the centroids of the seven clusters with the highest Global Explained Variance (GEV) for each subject.

Subsequently, to establish a standard reference for the microstate fitting process, the K-means algorithm was applied at the group level. To obtain topographical maps for the combined PRE+POST dataset, we clustered the dominant individual topographies for the PRE-VR-ES, PRE-VR-CTRL, POST-VR-ES and POST-VR-CTRL experimental conditions. Furthermore, we performed a group-level analysis of EEG signals recorded during the VR experience, which involved a separate clustering process using individual data solutions for each condition (VR-ES and VR-CTRL). The optimal number of clusters at the group level was determined using criteria implemented in Cartool v5.02 (an open-source academic software developed by Denis Brunet; cartoolcommunity.unige.ch), based on seven maximally independent criteria: Davies and Bouldin, Gamma, Silhouette, Dunn Robust, Point-Biserial, Krzanowski-Lai Index, and Cross-Validation (Brunet et al., 2011; Custo et al., 2017).

In the second part of the microstate analysis, all acquired and pre-processed EEG signals were subjected to the fitting process. We conducted the analysis separately for each condition (PRE-POST/ VR experience), applying the templates derived from the group-level clustering. A temporal smoothing (window half-size 3 (24 ms), Besag factor of 10, and a rejection of small-time frames (when < 3, i.e., 24 ms) were applied. To determine the temporal parameters of microstates, each time point in the individual data was assigned to the microstate cluster with which it correlated the best (Brunet et al., 2011). A correlation coefficient threshold of 0.6 was used to exclude short, transient noise frames in the EEG signal.

### Statistical Analysis

We evaluated whether data sets of microstate parameters, mean duration (ms), and occurrence (Hz) could be fitted to a Gaussian distribution. Using the Shapiro-Wilk normality test, we concluded that our data did not follow a normal distribution. Next, to explore VR-induced changes in the temporal dynamics of networks, we applied non-parametric Wilcoxon signed-rank two-tailed tests to compare PRE and POST results within each group (VR-ES, VR-CTRL) and a non-parametric Mann-Whitney independent-samples test to identify differences between groups during the VR experience. We adjusted p-values for multiple comparisons using the false discovery rate correction (FDR) (Benjamini, 1995). To quantify the size effect of the Wilcoxon signed-rank two-tailed tests or the Mann-Whitney independent-samples test, we calculated the biserial rank coefficient (rb) (Kerby, 2014). We assess the rb values according to McGrath and Meyer, whose interpretation is as follows: a limited statistical (small) effect if rb < 0.10, a medium if 0.10 ≤ rb ≤ 0.37, and an evident (high) statistical effect if rb > 0.37 (Fritz et al., 2012; McGrath & Meyer, 2006).

Changes over time in behavioural traits (emotional and anxiety reactivity) or cortisol data were evaluated using Mixed ANOVA, with Tukey-corrected post-hoc comparisons. Partial eta-squared (η^2^_P_) was reported to indicate effect sizes for the repeated-measures ANOVA tests. We interpreted it considering Cohen’s rules: small effect (η^2^_P_∼0.01), medium (η^2^_P_∼0.06), and large effect (η^2^_P_∼0.14) (Cohen, 1988).

Furthermore, we conducted a moderation analysis to determine whether microstate descriptors and behavioural states (emotional and anxiety reactivity) or cortisol data are related. In the moderation models, we used behavioural states (emotional and anxiety reactivity) or cortisol data and EEG microstate temporal parameters as both dependent and independent variables, in interaction with high (if score>mean+ 1 SD, SD-standard deviation), average or low (if score<mean-1SD) resilience or trauma scores.

## Results

### Behavioural results

To assess the emotional reactivity after the VR-ES exposure, as compared to the VR-CTRL condition, we applied a 2×5 ANOVA with Group (VR-CTRL, VR-ES) and Time (T_0_, T_1_, T_2_, T_3_, T_4_) as within-subjects factors. Regarding emotional reactivity (**Figure 2a**, **Table 1**), the findings revealed a significant main effect of time (F(4,310)=9.07; η^2^_P_=0.105, p<0.001) and interaction (F(4,310)=5.86; η^2^_P_=0.070, p<0.001), but no effect of group (F(1,310)=3.56; η^2^_P_=0.011, p=0.06). When we performed post hoc analyses on the Time x Group interaction, we observed significant results in the VR-ES group only for T_0_ vs T_1_ (p_tukey_ = 0.04), with no significant differences for T_0_ vs T_2_ (p_tukey_ = 0.92), T_0_ vs T_3_ (p_tukey_ = 0.78), or T_0_ vs T_4_ (p_tukey_ = 0.75). In contrast, for the VR-CTRL Group, emotional reactivity scores at baseline were specifically higher than scores at each subsequent time point (T_0_ vs T_1_: p_tukey_=0.023, T_0_ vs T_2_: p_tukey_=0.006, T_0_ vs T_3_: p_tukey_<0.001, T_0_ vs T_4_: p_tukey_=0.006).

**Figure 2.**
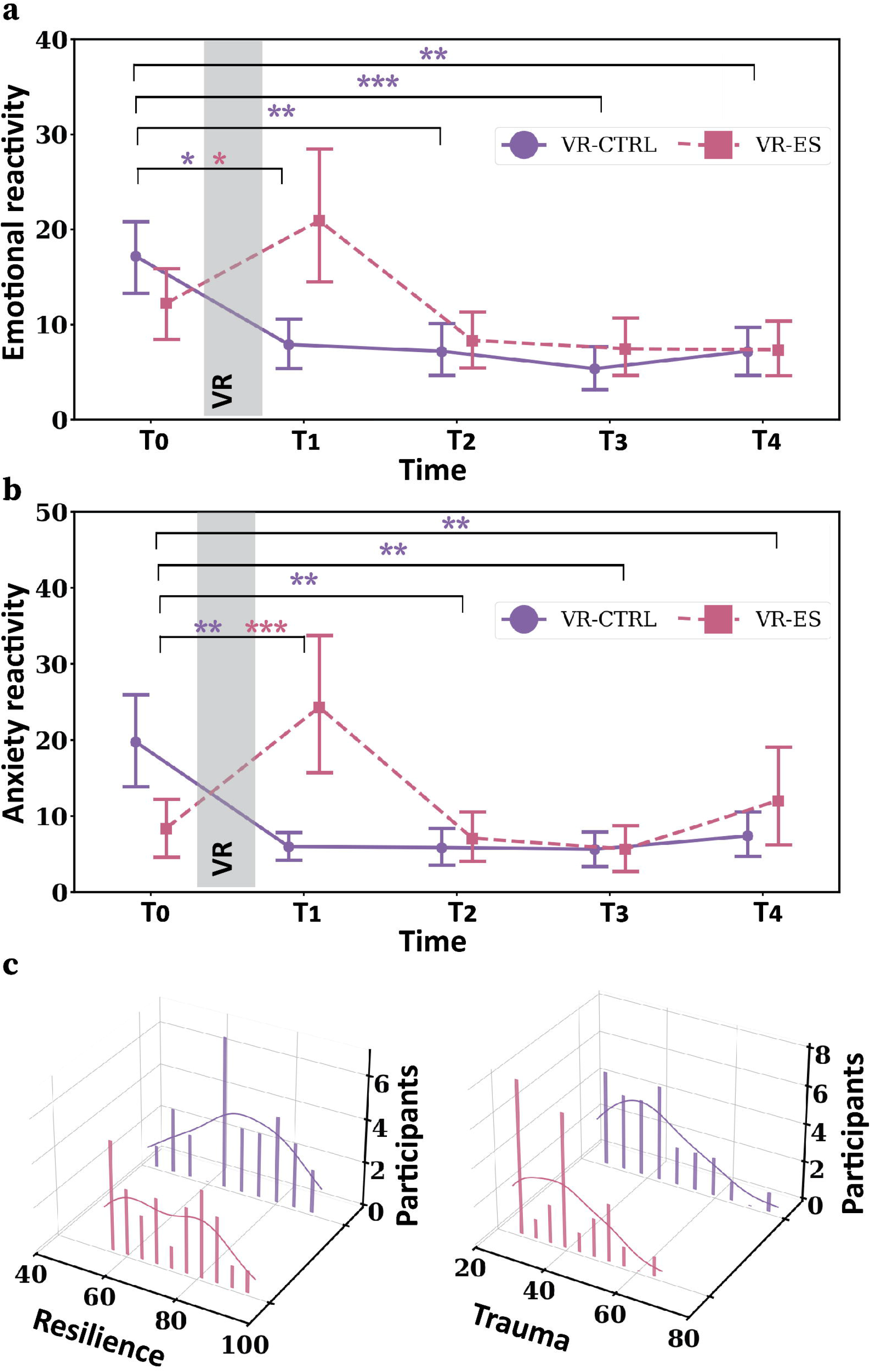
Behavioural states results. **(a, b)** Emotional and anxiety reactivity changes over time: scores over five collection time points (T_0_, T_1_, T_2_, T_3_, T_4_). The post hoc significant within-time differences are marked with Tukey-corrected p-values: *p<0.05, **p<0.01, ***p<0.001. Error bars show 95% confidence intervals. The purple circle and pink square represent the mean value of the data for that time point. **(c)** Distribution of resilience and trauma scores in the study sample.

We also assessed changes in anxiety reactivity (**Figure 2b**, **Table 1**). A 2×5 ANOVA with Group (VR-CTRL, VR-ES) and Time (T_0_, T_1_, T_2_, T_3_, T_4_) factors also demonstrated significant main effect of time (F(4,305)=5.88; η^2^_P_=0.072, p<0.001) and interaction (F(4,305)=9.20; η^2^_P_=0.108, p<0.001), but no effect of Group (F(1,305)=2.68; η^2^_P_=0.007, p=0.103). Performing Tukey-adjusted post hoc comparisons, we found a significant difference in the VR-ES group only at T_0_ vs T_1_ (p_tukey_<0.001). T_0_ vs T_2_ (p_tukey_=1.00), T_0_ vs T_3_ (p_tukey_=0.99), and T_0_ vs T_4_ (p_tukey_=0.99) did not show statistically significant changes. In contrast, in the VR-CTRL group, follow-up comparisons indicated that anxiety scores at baseline were significantly higher than those at each of the later time points (T_0_ vs T_1_: p_tukey_=0.004; T_0_ vs T_2_: p_tukey_=0.003; T_0_ vs T_3_: p_tukey_=0.002; T_0_ vs T_4_: p_tukey_=0.009).

**Table 1.**
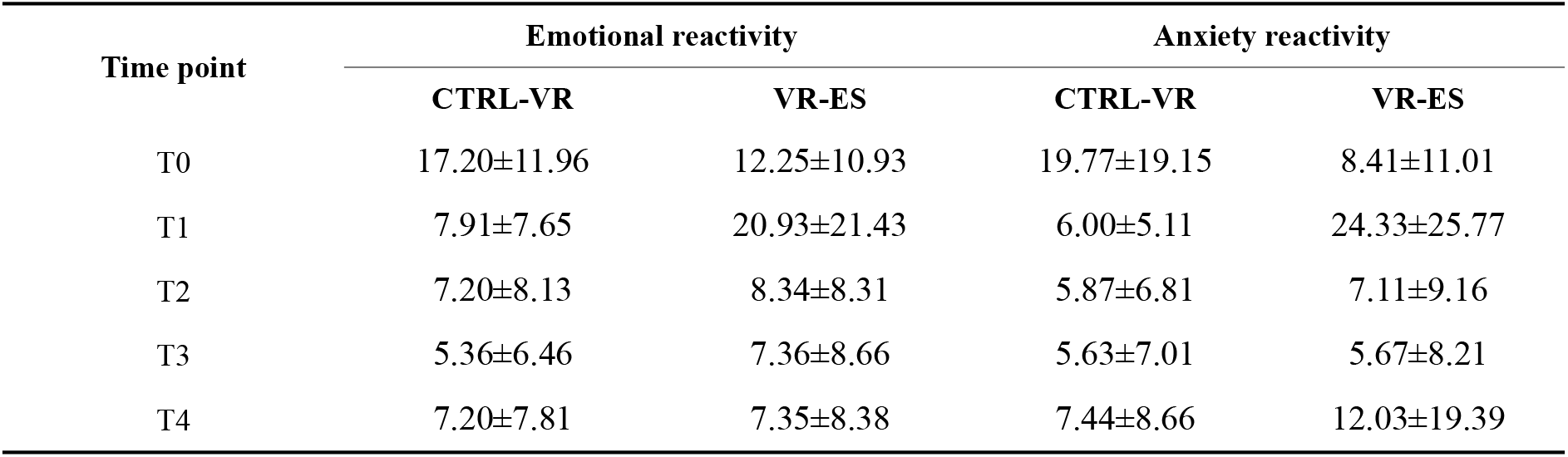
Emotional and anxiety reactivity scores (M ± SD) for each time point.

In **Figure 2c** we described resilience and trauma scores within our cohort.

### Hormonal results

To assess stress-related hormonal changes after VR-ES, we normalized the cortisol data, applying a logarithmic transformation (Miller & Plessow, 2013). Then, we performed a 2×5 ANOVA with Group (VR-CTRL, VR-ES) and Time (T_0_, T_1_, T_2_, T_3_, T_4_) as the within-subjects factors (**Figure 3a**, **Table 2**). The results showed statically significant main effects of time (F(4, 316)=5.179; η^2^_P_=0.062, p<0.001) and group (F(1,316)=4.002; η^2^_P_=0.013, p=0.046) and no interaction: F(4,316)=0.764; η^2^_P_=0.011, p=0.549). The Tukey-adjusted post-hoc comparisons for Time x Group did not indicate significant results in the VR-ES group (T_0_ vs T_1_: p_tukey_=0.945, T_0_ vs T_2_: p_tukey_=0.941, T_0_ vs T_3_: p_tukey_=0.913, T_0_ vs T_4_: p_tukey_=0.729). In contrast, in the VR-CTRL group, follow-up comparisons revealed that cortisol levels at T_0_ were significantly higher than those measured at T_2_, T_3_, and T_4_ (T_0_ vs T_1_: p_tukey_=0.292; T_0_ vs T_2_: p_tukey_=0.047; T_0_ vs T_3_: p_tukey_=0.012; T_0_ vs T_4_: p_tukey_=0.001).

**Figure 3.**
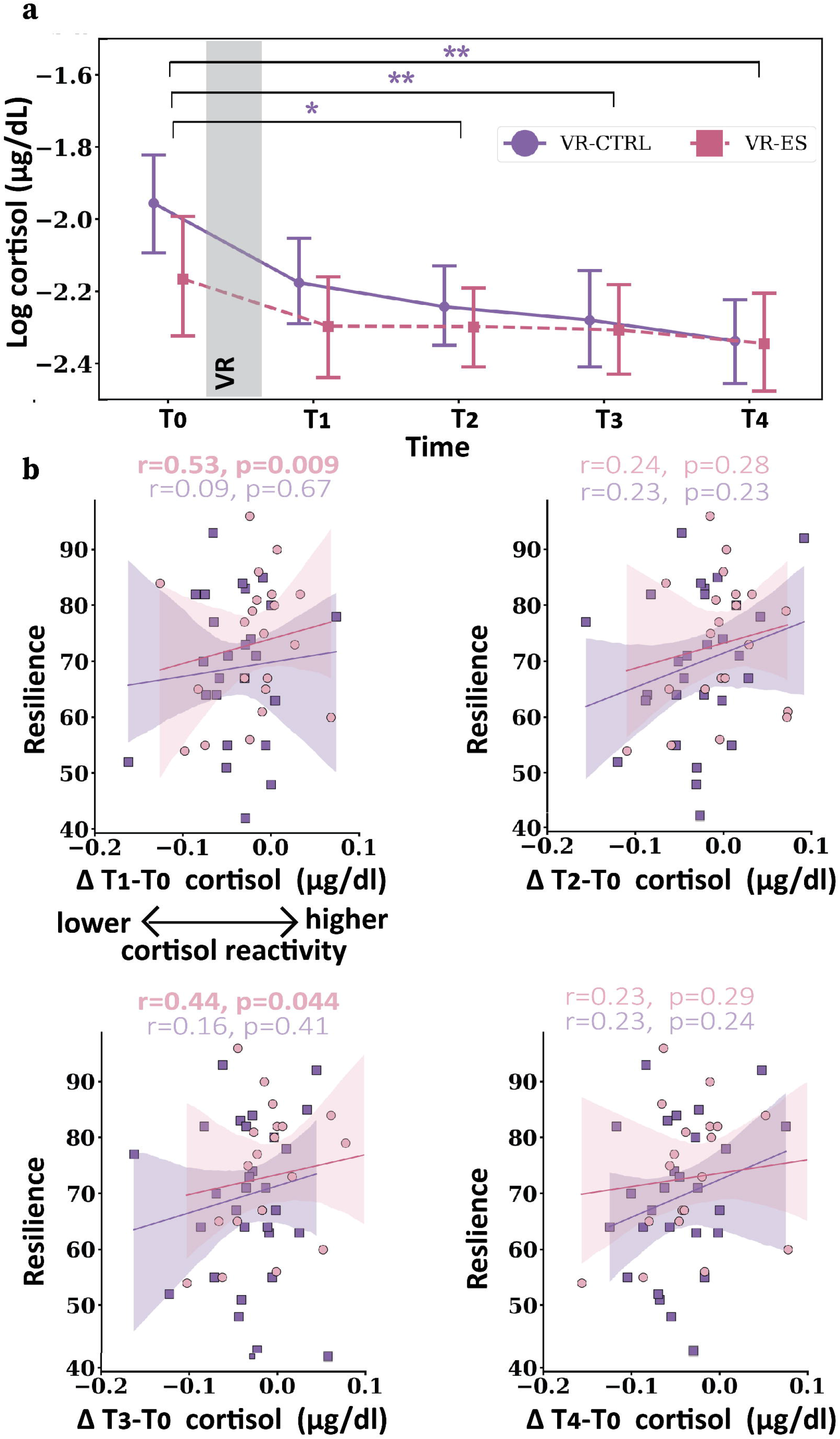
Cortisol results. **(a)** Log-transformed cortisol data over five collection time points (T_0_, T_1_, T_2_, T_3_, T_4_). The post hoc significant within-time differences are marked with Tukey-corrected p-values: *p<0.05, **p<0.01. Error bars show 95% confidence intervals. The purple circle and pink square represent the mean value of the data for that time point. **(b)** Spearman’s rank correlations between cortisol change scores and resilience. Lines are linear regression predictions; shaded areas are 95% confidence intervals.

**Table 2.** Cortisol levels (μg/dL) (M ± SD) for each time point.

| Time point | Cortisol ($\mu\text{g/dL}$) | | Log cortisol ($\mu\text{g/dL}$) | |
| --- | --- | --- | --- | --- |
|  | CTRL-VR | VR-ES | CTRL-VR | VR-ES |
| $T_0$ | $0.18 \pm 0.09$ | $0.14 \pm 0.12$ | $-1.84 \pm 0.48$ | $-2.11 \pm 0.54$ |
| $T_1$ | $0.13 \pm 0.08$ | $0.11 \pm 0.06$ | $-2.11 \pm 0.44$ | $-2.25 \pm 0.44$ |
| $T_2$ | $0.12 \pm 0.06$ | $0.11 \pm 0.04$ | $-2.19 \pm 0.51$ | $-2.26 \pm 0.38$ |
| $T_3$ | $0.12 \pm 0.06$ | $0.11 \pm 0.05$ | $-2.22 \pm 0.45$ | $-2.27 \pm 0.42$ |
| $T_4$ | $0.11 \pm 0.05$ | $0.10 \pm 0.05$ | $-2.32 \pm 0.43$ | $-2.31 \pm 0.44$ |

To assess individual variability in cortisol data as a function of self-reported resilience and adverse childhood experiences, we examined, for both experimental conditions, whether changes in cortisol levels over time were linked to resilience or trauma. Thus, we computed correlations between ΔT_i+1_-T_i_ cortisol (i=0,1,2,3) and the resilience and trauma score, as measured by CD-RISK and CTQ, respectively. In the VR-ES group, we found a significant positive relationship between change scores in cortisol levels and resilience at ΔT_1_-T_0_ (r=0.53, p=0.009) and ΔT_3_-T_0_ (r=0.44, p=0.044). Although the correlations for ΔT_2_-T_0_ cortisol (r=0.24, p=0.28) and ΔT_4_-T_0_ cortisol (r=0.23, p=0.29) were not statistically significant, they showed the same positive trend, aligning with the pattern observed for ΔT_1_–T_0_ cortisol and ΔT_3_–T_0_ cortisol. In the VR-CTRL group, cortisol change scores were not significantly correlated with resilience at any time point (**Figure 3b**). Lastly, in neither group did the correlation analyses reveal significant associations between cortisol change scores and trauma. Thus, only in the VR-ES condition did self-perceived resilience predict cortisol changes, with greater perceived resilience associated with increased cortisol release.

### Topographical results

Using seven independent criteria (detailed in Methods), we objectively determined that six microstates best describe the variability in the combined PRE and POST dataset (before and after VR-CTRL/VR-ES), accounting for 90.41% of the explained variance (**Figure 4a**). In addition, analysis of EEG data recorded during the VR experience showed that the cluster algorithm robustly identified six topographical microstates, explaining 84.57% of the variance in the VR-CTRL group and 85.14% in the VR-ES group (**Figure 4b**). These were ordered A to F based on their similarity to the literature (Custo et al., 2017; Michel & Koenig, 2018; Tarailis et al., 2024). Moreover, when comparing topographies between VR-CTRL broadband and VR-ES broadband (i.e., the diagonal entries in the correlation matrix), all spatial correlations showed r > 0.90 (**Figure 4c**), supporting the validity of comparing microstate temporal parameters across experimental conditions.

**Figure 4.**
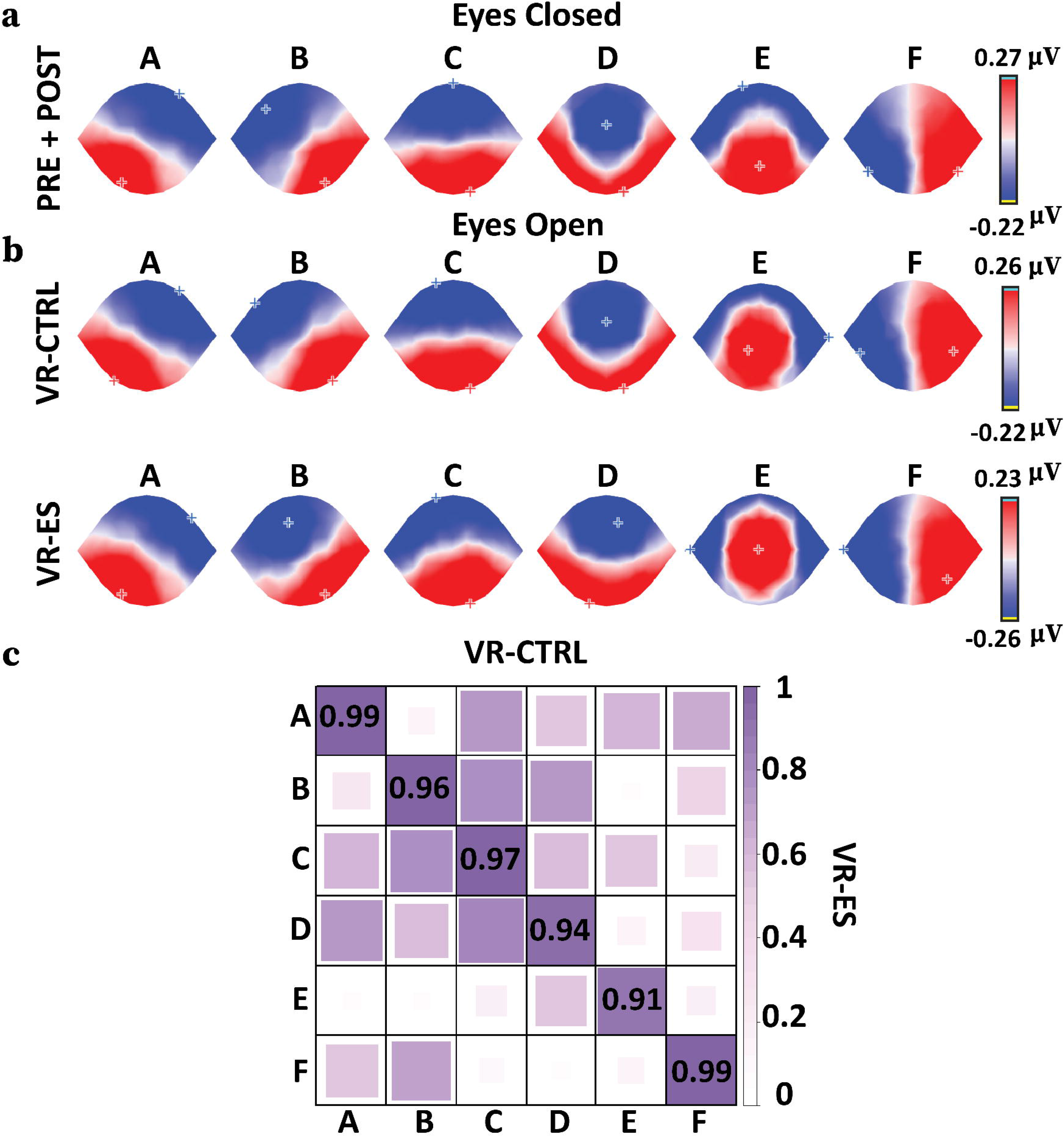
Microstates spatial distribution. **(a)** Topographical maps A to F for PRE+POST (VR-CTRL+VR-ES) conditions (red: positive amplitudes, blue: negative amplitudes). MS A exhibits a left-right orientation, MS B presents a right-left orientation, MS C electric activity varies from frontal to occipital, MS D covers from centro-frontal to lateral and occipital, and from centro-occipital to frontal in MS E and MS F, with a left-lateralized maximum. **(b)** Identified classes of microstates for VR-CTRL and VR-ES groups. **(c)** Spatial correlation results for microstate topographies under VR-CTRL and VR-ES conditions (*r* values).

### Inter-group differences in neural activity between the VR-ES and VR-CTRL during freely moving explorations

To assess the EEG microstate temporal dynamics during VR-ES as compared to VR-CTRL, we applied independent-samples t-tests. Regarding mean duration (**Figure 5a**), MS C lasted for a shorter amount of time in the VR-ES group compared to the VR-CTRL group (VR-CTRL: 94.09±12.21, VR-ES: 83.78±10.41, p<0.001, r_b_=0.55, FDR corrected). In contrast, MS D showed an opposite pattern (VR-CTRL: 76.89±6.34, VR-ES: 87.51±12.03, p<0.001, r_b_=0.52, FDR corrected). Additionally, MS C (VR-CTRL: 2.59±0.41, VR-ES: 2.24±0.38, p=0.003, r_b_=0.46, FDR corrected) and MS B (VR-CTRL: 1.97±0.56, VR-ES: 1.68±0.54, p=0.045, r_b_=0.33, FDR corrected) occur less frequently in the VR-ES than the VR-CTRL group, whereas MS D showed a significantly higher occurrence in the VR-ES compared to the VR-CTRL group (VR-CTRL: 1.71±0.54, VR-ES: 2.42±0.62, p<0.001, r_b_=0.63, FDR corrected) (**Figure 5b**). We did not find significant differences for MS A, E, and F.

**Figure 5.**
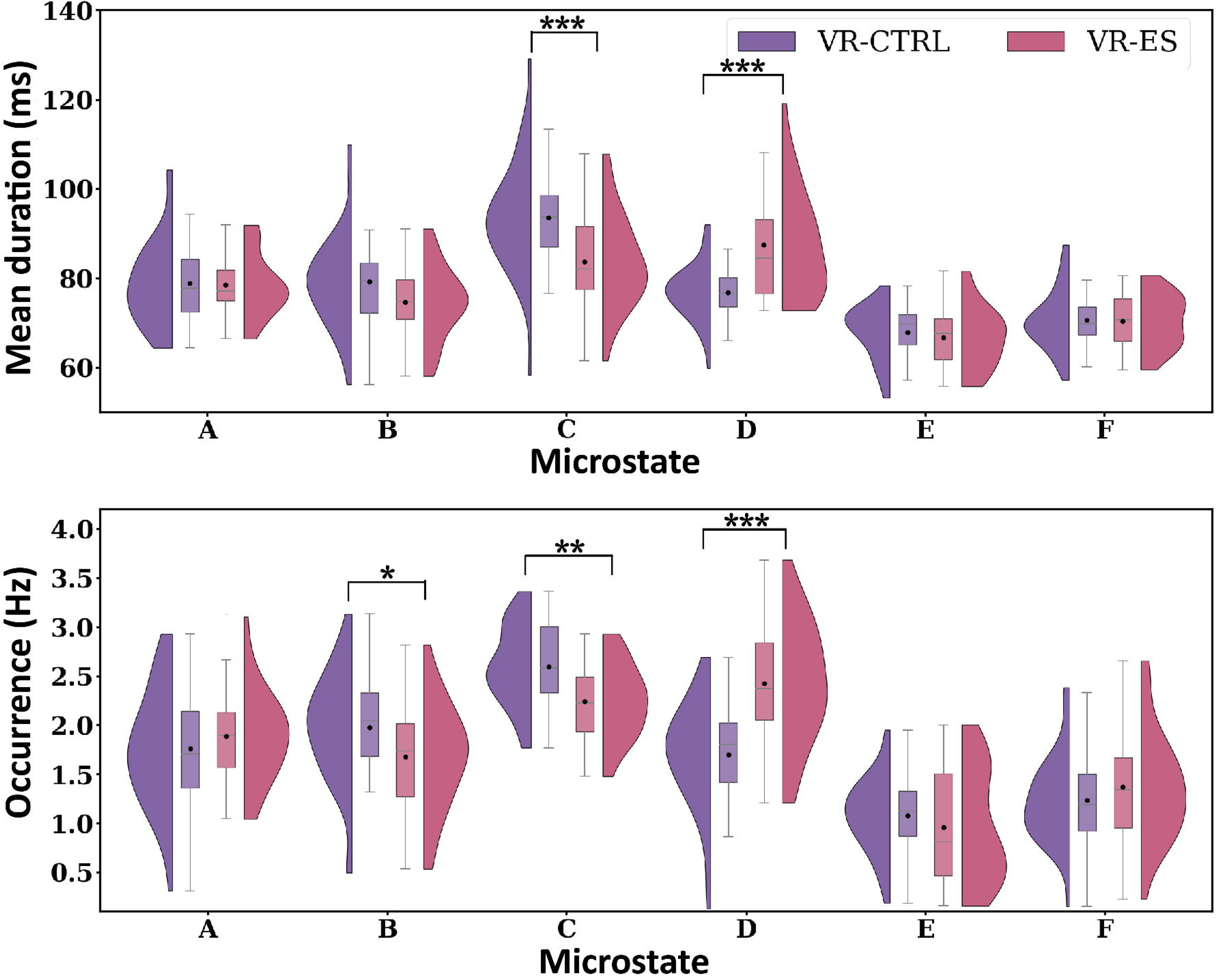
VR experience modulates microstate temporal dynamics: during VR results. **(a)** Microstate mean duration changes during VR-ES. **(b)** Microstate occurrence changes over time. The significant intergroup differences are marked with FDR-corrected p-values: *p<0.05, **p<0.01, ***p<0.001. In the boxplots, the lower and upper boundaries correspond to the first (Q1) and third (Q3) quartiles, respectively, while the middle line represents the median. The ends of the vertical grey lines indicate the values corresponding to Q1-1.5IQR and Q3-1.5IQR, respectively (IQR-interquartile range). The black circle is the mean value of the data.

### Moderated associations between MS temporal dynamics during VR-ES and recovery of behavioural, emotional, and hormonal states

We conducted moderation analyses to investigate whether microstate temporal-dynamic descriptors (i.e., mean duration and occurrence) extracted from EEG recorded during VR experience are associated with behavioural and hormonal states (emotional and anxiety reactivity or cortisol data).

For the MS C mean duration VR-ES changes, we found a significant moderation of resilience traits on the associations with emotional reactivity. Decreased mean duration of MS C during VR-ES is positively associated with lower emotional reactivity (ΔT_1_-T_0_) in high-resilience (+1 SD) individuals (**Figure 6a**, **Table 3**) and with decreased (ΔT_1_-T_0_) anxiety levels in low-trauma (-1 SD) participants (**Figure 6c**, **Table 3**). Also, we identified a negative relation between VR-ES MS C occurrence and ΔT_2_-T_1_ cortisol change scores in low trauma individuals (**Figure 6e**, **Table 3**). Decreased MS C occurrence during VR-ES predicts a higher cortisol change score ΔT_2_-T_1_ in low-trauma individuals after VR-ES.

**Figure 6.**
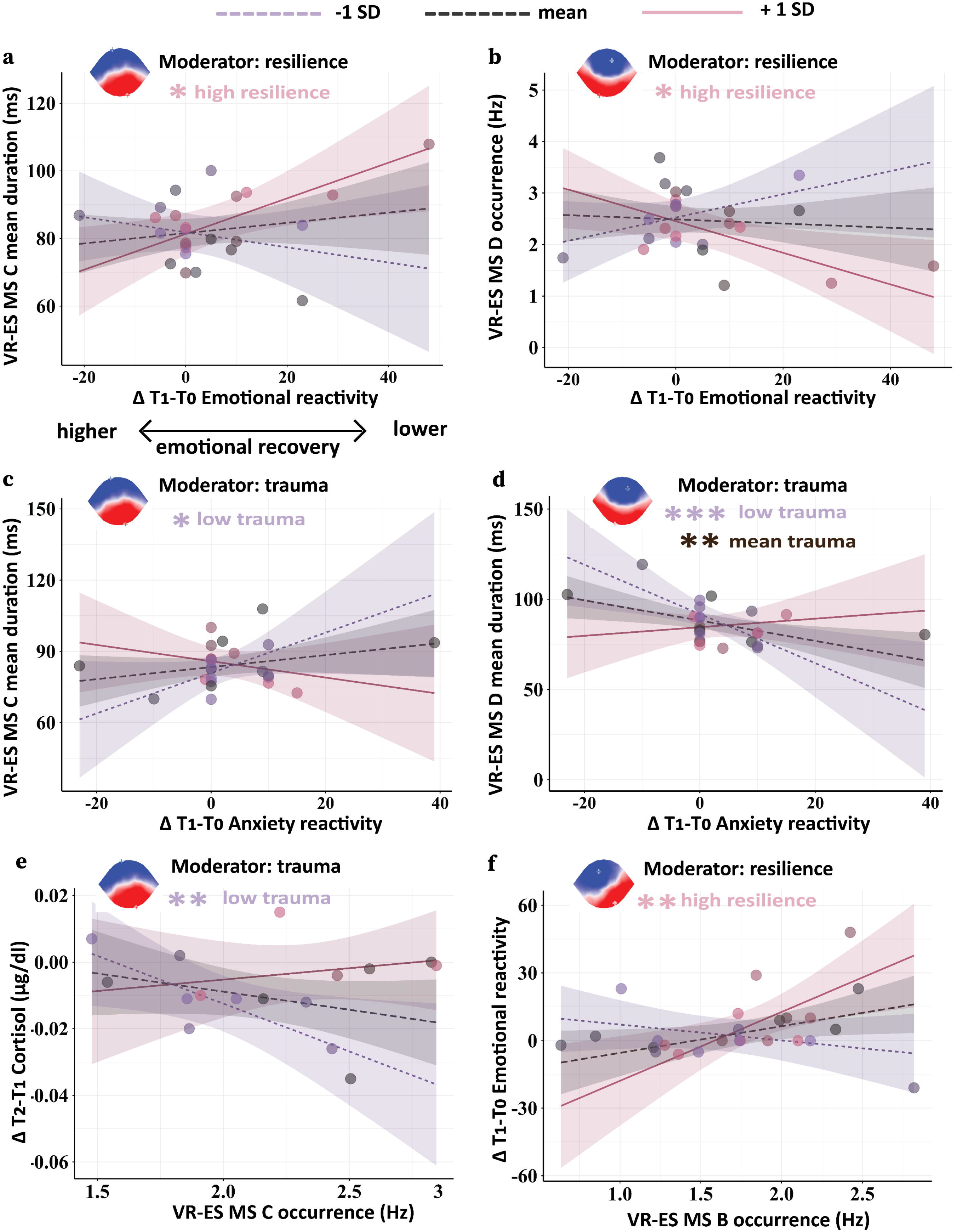
Resilience and trauma moderate changes in the temporal dynamics of microstates: during VR results. **(a, b)** Resilience moderates the association between VR-ES MS C mean duration & VR-ES MS D occurrence and ΔT1-T0 emotional reactivity change score. **(c, d)** Trauma moderates the association between VR-ES MS C mean duration & VR-ES MS D mean duration and ΔT1-T0 emotional reactivity change score. **(e)** Trauma moderates the association between ΔT2-T1 cortisol change score and VR-ES MS C occurrence. **(f)** Resilience moderates the association between ΔT1-T0 emotional reactivity change score and VR-ES MS B occurrence. Purple dots and regression lines represent individuals with one standard deviation below the mean (−1 SD) on moderator distribution. Pink dots and regression lines correspond to individuals with one standard deviation above the mean (+1 SD) on the moderator. Black dots and regression lines depict the individuals whose moderator score falls between one standard deviation below and one standard deviation above the mean. The significant slope differences (-1 SD, mean, +1 SD) are marked with p-values: *p<0.05, **p<0.01, ***p<0.001.

**Table 3.**
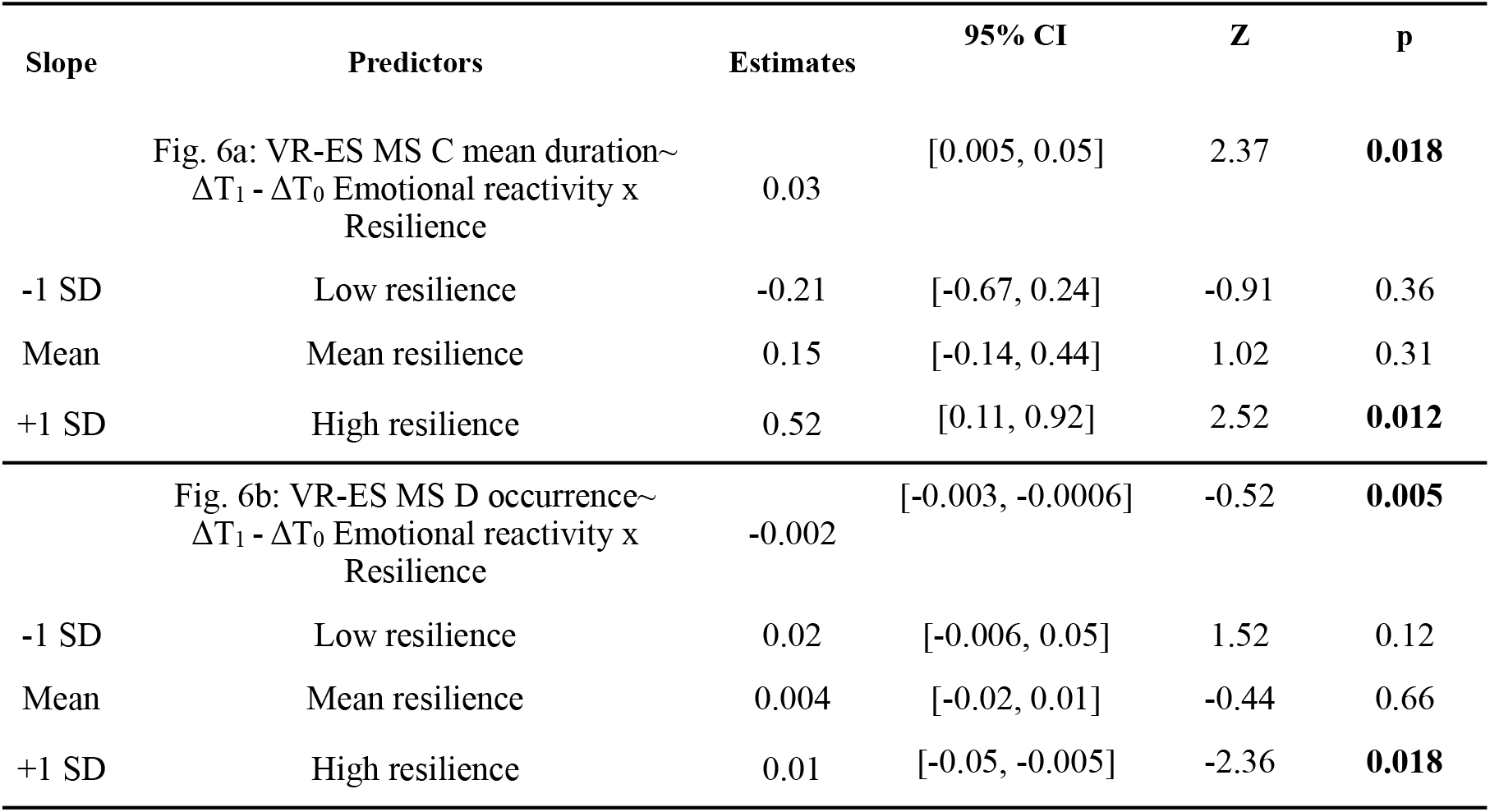
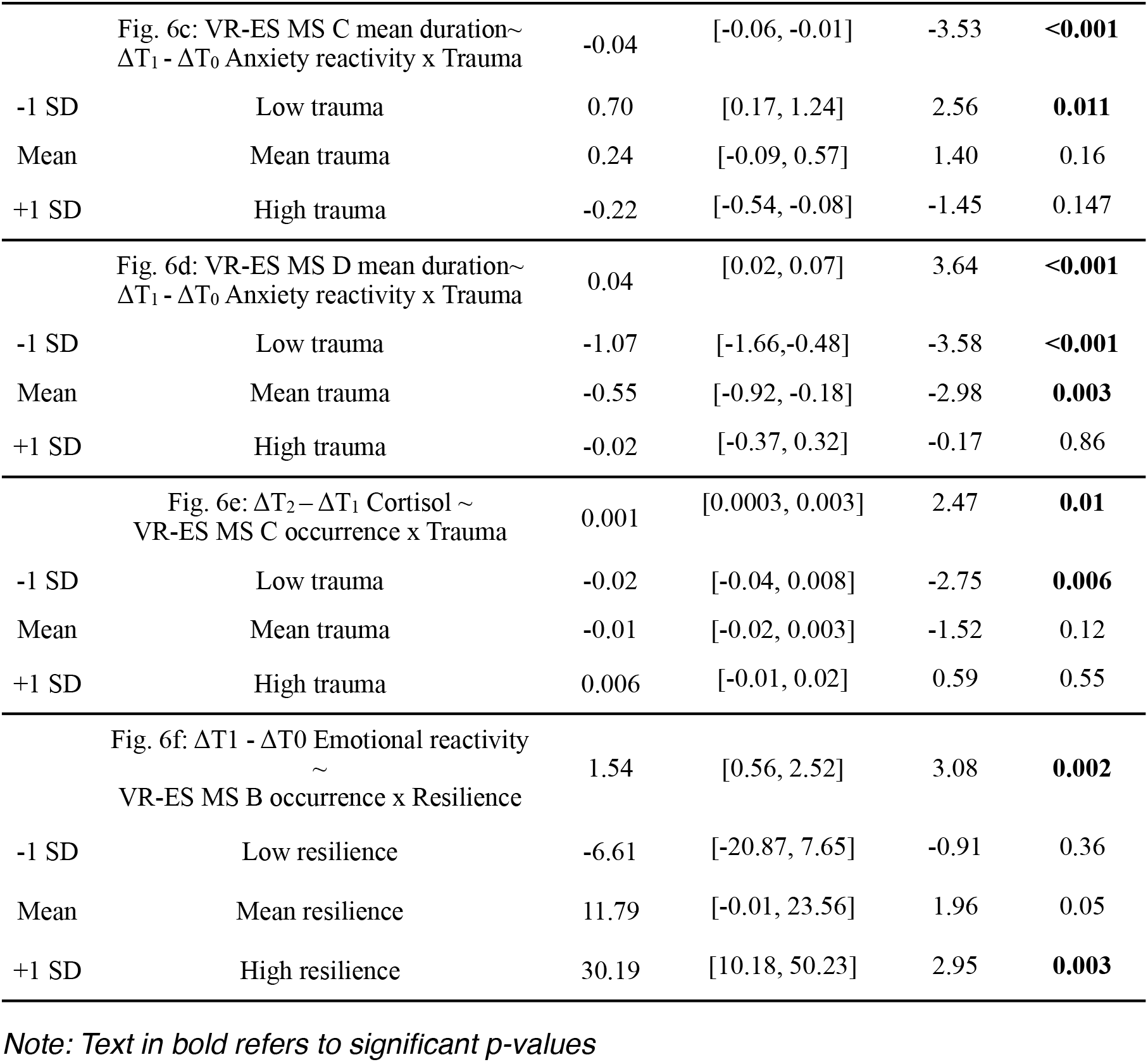
Moderation analysis results for B, C, and D microstates & behavioural traits/ cortisol.

Additionally, the relationship between emotional reactivity/anxiety for MS D occurrence was significantly moderated by resilience traits and childhood trauma. We observed a significant negative relation between MS D occurrence and ΔT_1_-T_0_ emotional reactivity in high resilience (+1 SD) individuals (**Figure 6b**, **Table 3**) and between MS D mean duration and ΔT_1_-T_0_ anxiety in low (- 1 SD) and mean trauma participants (**Figure 6d**, **Table 3**). Increased MS D temporal dynamics during VR-ES predict lower emotional reactivity and anxiety in high-resilience and low-trauma individuals after VR-ES.

Finally, MS B showed a specifically positive association with ΔT_1_-T_0_ emotional reactivity in high-resilience individuals (**Figure 6f**, **Table 3**).

### Intra-group differences across time in neural activity during recovery after VR- ES and VR-CTRL

To investigate PRE-POST alterations in microstate duration (ms) and frequency (Hz) induced by the VR-ES and VR-CTRL experiences, we performed two-tailed Wilcoxon signed-rank tests comparing POST to PRE measurements.

In terms of mean duration (**Figure 7c**), we found a significant decrease of MS E (PRE VR-ES: 75.44±6.06, POST VR-ES: 73.51±6.31, p=0.042, r_b_=0.58, FDR corrected). Concerning the occurrence of MSs (**Figure 7d**), MS C occurs more frequently (PRE VR-ES: 2.25±0.31, POST VR-ES: 2.35±0.33, p=0.044, r_b_=0.55, FDR corrected), while MS D exhibited a significantly reduced occurrence rate (PRE VR-ES: 1.85±0.52, POST VR-ES: 1.66±0.59, p=0.042, r_b_=0.59, FDR corrected). In the VR-ES group, MS A, B and F did not reveal significant differences in mean duration or occurrence after the VR experience.

**Figure 7.**
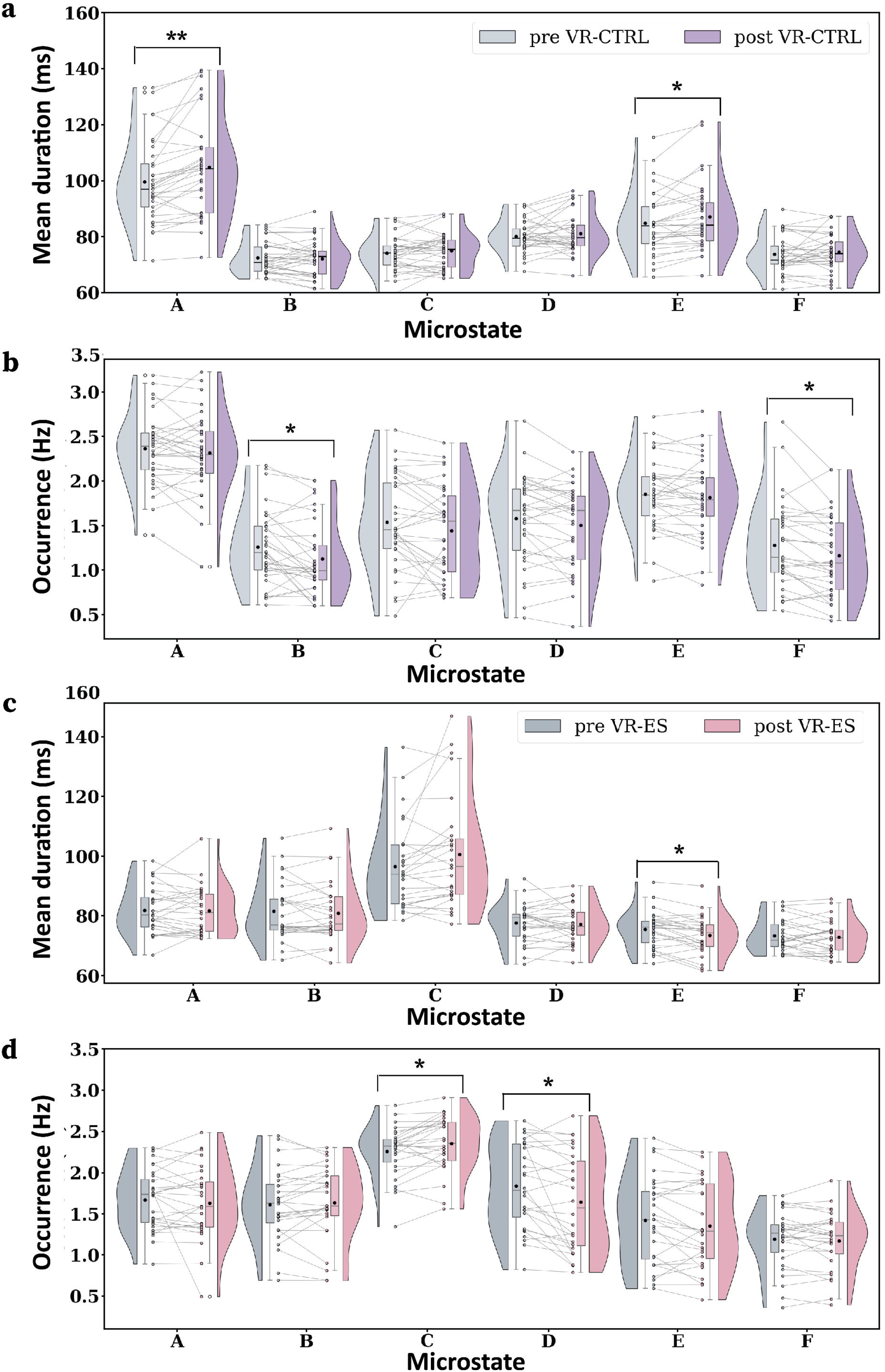
VR experience modulates microstate temporal dynamics: pre vs post results. **Changes in mean duration and occurrence for each class of microstate. (a, b)** VR-CTRL group: **(c, d)** VR-ES group. The significant inter-session differences are marked with FDR corrected p-values *p<0.05,**p<0.01. In the boxplots, the lower and upper boundaries correspond to the first (Q1) and third (Q3) quartiles, respectively, while the middle line represents the median. The ends of the vertical grey lines indicate the values corresponding to Q1-1.5IQR and Q3-1.5IQR, respectively. The black circle represents the mean of the data.

In the VR-CTRL group, the effects for MS C and D were not statistically significant. After the VR experience, two classes of MSs exhibited a significant increase in duration (**Figure 7a**): MS A (PRE VR-CTRL: 99.71±14.51, POST VR-CTRL: 104.83±17.61, p=0.0012, r_b_=0.72, FDR corrected) and MS E (PRE VR-CTRL: 84.83±11.03, POST VR-CTRL: 87.12±12.35, p=0.024, r_b_=0.55, FDR corrected). Regarding the occurrence (**Figure 7b**), we found that MS B (PRE VR-CTRL: 1.25±0.41, POST VR-CTRL: 1.12±0.42, p=0.024, r_b_=0.53, FDR corrected) and MS F (PRE VR-CTRL: 1.27±0.48, POST VR-CTRL: 1.13±0.41, p=0.024, r_b_=0.53, FDR corrected) showed decreased occurrences.

### Moderated associations between the MS temporal dynamics during VR-ES recovery and behavioural states

We analysed whether the differences between behavioural states and MS microstate temporal-dynamic descriptors during recovery (i.e., change scores for mean duration and occurrence) are moderated by resilience traits and/or childhood trauma. Applying a moderation analysis, we found that Δ POST-PRE VR-ES MS C occurrence relation with emotional reactivity is moderated by childhood trauma only. We found a positive association between MS D occurrence and the ΔT_1_-T_0_ emotional reactivity change score only in low-trauma (-1 SD) individuals (**Figure 8a**, **Table 4**). Decreased MS D mean occurrence during recovery phase predicts a lower emotional reactivity change score ΔT_1_-T_0_ in low-trauma individuals.

**Figure 8.**
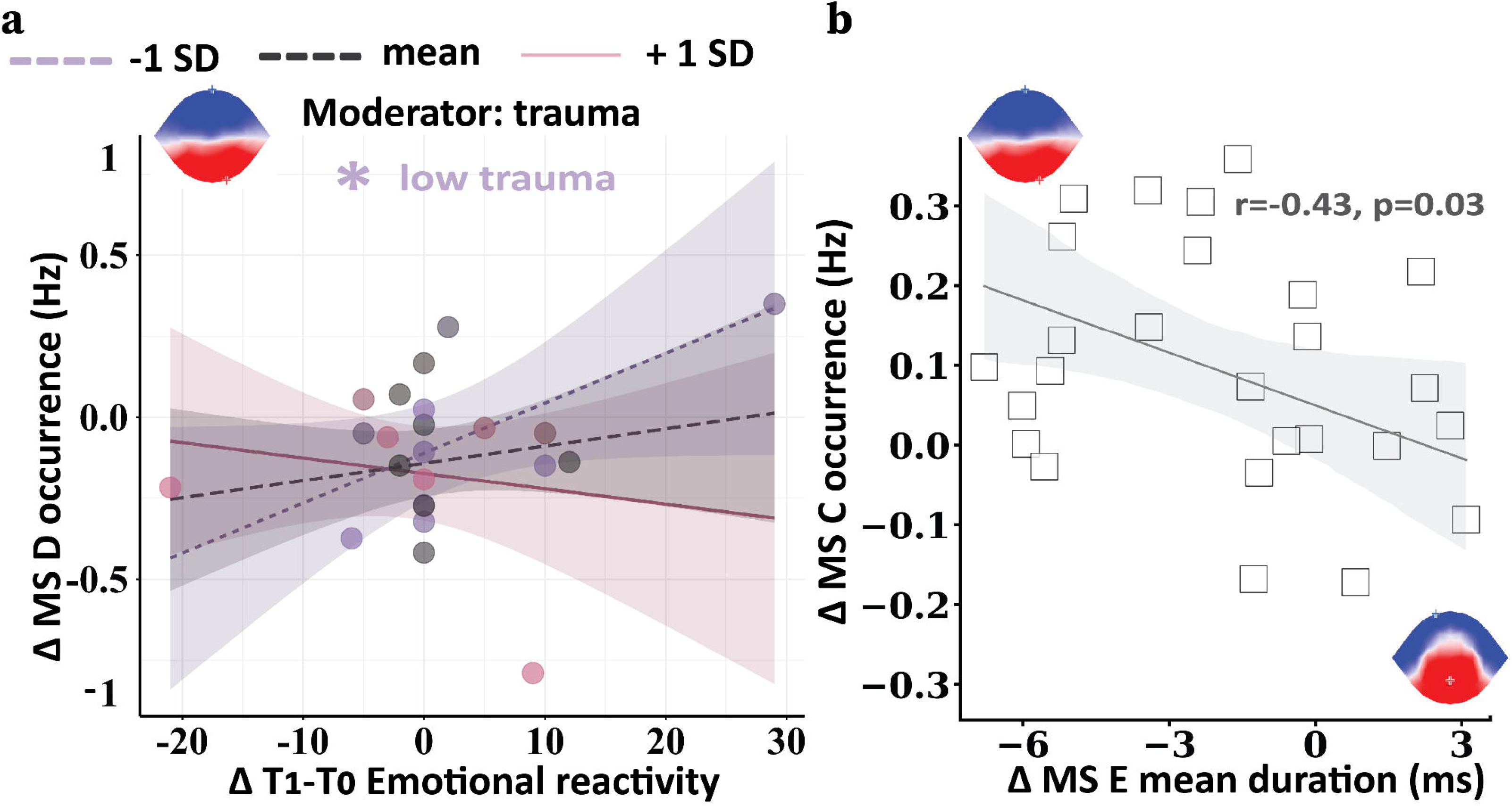
Microstates temporal dynamics change scores (Δ post-pre): Moderation and Correlation. **(a)** Trauma moderates the association between Δ POST-PRE VR-ES MS D occurrence and ΔT1-T0 emotional reactivity change score. Purple dots and regression lines denote individuals with one standard deviation below the mean (−1 SD) on trauma distribution. Pink dots and regression lines correspond to individuals with one standard deviation above the mean (+1 SD) of trauma scores. Black dots and regression lines are associated with individuals whose trauma scores fall between one standard deviation below and one standard deviation above the mean. The significant slope analysis differences (-1 SD, mean, +1 SD) are marked with p-values *p<0.05. **(b)** Spearman rank correlation between Δ POST-PRE VR-ES MS C occurrence and Δ POST-PRE VR-ES MS E mean duration. Lines depict linear regression predictions; shaded areas are 95% Confidence intervals.

**Table 4.** Moderation analysis results for D microstate & Emotional reactivity.

| Slope | Predictors | Estimates | 95% CI | Z | p |
| --- | --- | --- | --- | --- | --- |
| | Fig. 8a: ΔPOST-PRE VR-ES MS D occurrence rate~<br>$\Delta T_1 - \Delta T_0$ Emotional reactivity x Trauma | -0.0009 | [-0.001, -0.00006] | -2.11 | **0.036** |
| -1 SD | Low trauma | 0.01 | [0.0007, 0.03] | 2.06 | **0.039** |
| Mean | Mean trauma | 0.005 | [-0.005, 0.01] | 0.99 | 0.32 |
| +1 SD | High trauma | -0.004 | [-0.01, 0.009] | -0.63 | 0.52 |

*Note: Text in bold refers to significant p-values*

In addition, we found that change scores in MS C occurrence and MS E mean duration were significantly correlated (r=-0.43, p=0.03) (**Figure 8b**). The more Δ POST-PRE VR-ES MS C occurrence increases, the more Δ POST-PRE VR-ES MS E mean duration decreases.

### Exploratory MS analysis during VR-ES and recovery in cortisol responder participants

Possibly due to the relatively small sample size, we did not detect a significant moderating effect of cortisol. To further investigate if cortisol modulations might have an effect on the MS temporal dynamics, we divided our group into two groups based on the comparison between ΔT1-T0 and ΔT2-T1.

In subjects with ΔT_2_-T_1_ > ΔT_1_-T_0,_ cortisol (classified as responders) showed that, during the VR-ES experience, responders exhibited a significant decrease (p=0.011, r_b_=0.59, Mann-Whitney test) in the occurrence of MS C (responders: 2.07±0.34; non-responders: 2.42±0.31). In non-responder individuals, we did not find effects of cortisol modulations on the VR-ES MS temporal dynamics (**Figure 9a**).

**Figure 9.**
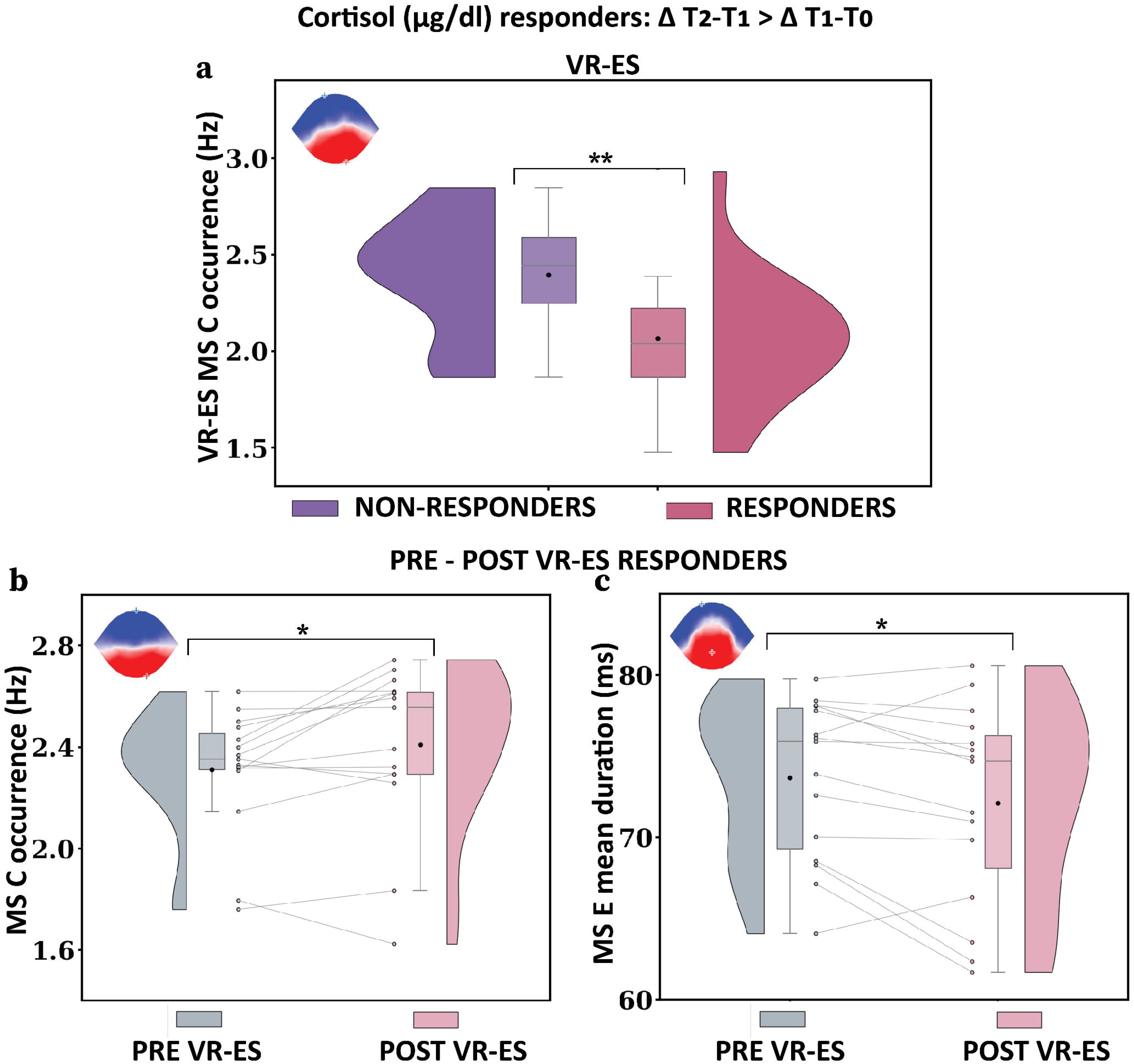
Microstate temporal dynamics in relation to cortisol changes. **(a)** Changes in VR-ES MS C occurrence in responder individuals vs non-responder participants. The significant intergroup differences are marked with ** (p<0.01). **(b)** Changes in MS C occurrence: pre-VR-ES responders vs. post-VR-ES responders. **(c)** Changes in MS E mean duration: pre VR-ES responders vs post VR-ES responders. The significant inter-session differences are marked with * (p<0.05).

When comparing PRE- and POST-intervention measures, in cortisol responders, MS C occurred more frequently (p=0.035, rb=0.62, two-tailed Wilcoxon test) in post VR-ES (2.42±0.32) than in pre-VR-ES (2.31±0.08) (**Figure 9b**). At the same time, MS E showed a significantly lower mean duration (p=0.035, r_b_=0.62, Wilcoxon matched-pairs signed rank two-tailed test) in POST VR-ES (72.11±6.18) compared to PRE VR-ES (73.67±4.91). We did not observe any significant changes in MS dynamics in non-responding participants (**Figure 9c**).

## Discussion

Natural disasters expose how rapidly the environment changes and how humans struggle to adapt, underscoring the need to understand the neural mechanisms underlying rapid reactivity and recovery. Although much research has focused on the positive effects of exposure to nature on human well-being and mental health and their neuronal correlates (e.g., Daube et al., 2026), less is known about how humans rapidly adapt to rising environmental challenges. Here, by combining multivariate analyses of prior adverse experiences during childhood, resilient psychological traits, hormonal reactivity, momentary affective states, and mobile EEG during an immersive VR earthquake experience, this study examines how large-scale brain dynamics adapt to environmental threat and support rapid emotional and physiological recovery as a function of resilience and prior exposure to adversity.

### Summary of findings

Earthquake VR-ES exposure significantly alters affective, hormonal, and EEG activity as a function of resilience traits and adverse childhood experiences. In contrast to VR-CTRL, VR-ES led to a significant increase in emotional and anxiety reactivity and to sustained cortisol release, with more pronounced cortisol reactivity in more resilient individuals (**Figure 2** and **Figure 3**).

During recovery, after VR-ES experience, both emotional reactivity and anxiety scores increased significantly and remained higher than those of the VR-CTRL group (**Figure 2**). In contrast, the VR-CTRL group showed lower scores at T_1_, which remained relatively stable through T_4_.

After the VR session, the ES group experienced a slight, nonsignificant drop in salivary cortisol from T_0_ to T_1_, with levels establishing and becoming similar to the VR-CTRL group by T_3_ and T_4_ (**Figure 3a**). In contrast, the VR-CTRL group showed a continuous decrease in cortisol from T_0_ to T_4_.

In opposition to the VR-ES MS C and MS D dynamics during VR exposure, in the recovery stage, MS C increased, and MS D decreased in the resting-state after VR-ES (**Figure 7cd**). Also, MS E showed a reduction after VR-ES. Moreover, these changes were associated with emotional reactivity and were moderated by low trauma scores (**Figure 8a**).

Importantly, the pattern of spontaneous microstates is differentially modulated during and after the aversive experience, with high specificity for different stages of affective processing. More specifically, we found specific anticorrelated modulations of MS C and MS D temporal dynamics during VR-ES exposure with high effect sizes (**Figure 5**). During VR-ES, the dynamics of MSs B and C decreased, whereas the dynamics of MS D were increased when compared to VR-CTRL. Moreover, these modifications predicted emotional and anxiety reactivity as a function of resilience traits and childhood aversive experiences, where in highly resilient individuals, MS C showed a positive association (**Figure 6ac**), whereas MS D exhibited a negative association with emotional recovery (**Figure 6bd**). Additionally, aversive experiences moderate the MS C to cortisol recovery, where individuals with low trauma experiences reveal a negative relation between cortisol level and MS C (**Figure 6e**). In summary, EEG microstate patterns for highly resilient individuals reveal that the more the MS C decreased during the VR-ES and the more MS C increased during the POST-VR-ES resting-state, the faster the emotional and anxiety state regulate. While for the MS D, the opposite pattern was significant, more MS D during VR-ES and less POST VR-ES resting state (**Figure 7**), the faster the emotional and anxiety state regulation.

In addition, cortisol may contribute to the resilient pattern of neural reactivity, as we found in an exploratory analysis that cortisol responders (i.e., ΔT_2_-T_1_ > ΔT_1_-T_0_) showed reduced MS C during VR-ES exposure (**Figure 9a**), whereas during recovery, the MS C increased, and MS E decreased (**Figure 9bc**).

In the following paragraphs, we will discuss in detail the EEG microstate results obtained during the reactivity phase of the natural disaster exposure during VR-ES and the recovery phase of the POST VR-ES resting state activity.

### EEG MS modulations during VR-ES reactivity

Here, we present an in-depth discussion of the VR-ES MSs modulations observed during the reactivity phase of the natural disaster simulation in the VR-ES. MS C showed a decrease in mean duration and occurrence rate during VR-ES free movement, compared to VR-CTRL (**Figure 5**). Other studies have reported modulation of MSs during emotional reactivity in socio-emotional contexts, with a clear link to its role in emotion reactivity and regulation (see Schiller et al., 2024). Most importantly, Nazare and Tomescu contribute to the body of research showing a significantly reduced presence of MS C in self-generated negative arousal affective states (Nazare & Tomescu, 2024). In that study, participants selected autobiographical memories with negative or positive valence, thereby limiting the generalizability of the interpretations to internal, self-generated affect alone. Here, we extend these results as we observed a decrease in MS C also during VR-based exposure to natural disaster scenarios, suggesting that these patterns of modulations are not specific to the type of affective generation self or environmentally generated in VR settings, and are reflecting a basic mechanism of adaptation to aversiveness that predicts emotional regulation as a function of psychological resilience traits, and prior aversive experiences as suggested by our moderation analyses (**Figure 6ac**).

MS C has also been shown to decrease when participants engage in tasks with cognitive load higher than that at rest, such as arithmetic tasks (Kim et al., 2021; Seitzman et al., 2017; Tarailis et al., 2024). Functionally, MS C has been related to task–negative frontal–parietal regions (i.e., the posterior cingulate cortex-PCC, precuneus, and left angular gyrus), important hubs of the default mode network (DMN). Alternatively, when only the four canonical microstates are represented in the data, studies have reported that MS C is also partially associated with task-positive salience (Britz et al., 2010; Custo et al., 2017; Tarailis et al., 2024). Our results appear to support the first view, in which MS C behaves more like a task-negative MS, decreasing activity during highly emotionally engaging tasks. These results suggest that DMN deactivation during VR-ES may be adaptive, reducing task-negative thoughts, mind-wandering, and self-related thoughts to facilitate task engagement. Moreover, Satpute and Lindquist argue that while earlier work linked the DMN to general affective dimensions such as arousal and valence via areas like the ventromedial prefrontal cortex, their central contribution is to show that DMN engagement-especially in the posterior cingulate-is better understood as supporting conceptual abstraction across emotional instances, and integration of self-related experiences rather than reflecting autonomic arousal intensity per se (Satpute & Lindquist, 2019). While this may not have been the case during the VR-ES, another possibility is that, during the recovery (POST VR-ES), resting-state MS C might be associated with the experiential integration part of the self-referential activity. However, both of these phases of adaptation, reactivity during VR-ES and recovery during the POST-VR-ES, are associated with better affective recovery, suggesting that both deactivation and activation of the MS C DMN-related activity support emotional regulation in highly resilient individuals at different stages of adaptation.

In addition, to better understand the MS C decrease and the mechanisms of resilient adaptations, and to complement self-reports of resilient traits moderations, we examined physiological stress responses, focusing on salivary cortisol as an objective marker of HPA-axis activation; however, it is also a biological marker of resilience. Several reviews argue that cortisol is not merely a stress hormone but, when dynamically and context-appropriately regulated, serves as a physiological marker of adaptive stress regulation and resilience (de Kloet & Joëls, 2024)(de Kloet & Joëls, 2024). With this view in mind, not only did the resilience traits moderate the MS C to emotional reactivity associations, but we also found that VR-ES MS C occurrence predicts the slower salivary cortisol drop (ΔT_2_-T_1_) in individuals with low aversive childhood experiences after VR-ES. From this perspective, given the MS C association with DMN, our results are consistent with the functional Magnetic Resonance Imaging (fMRI) literature on the link between acute stress and DMN deactivation. Indeed, studies have shown that acute stress-induced cortisol responses are associated with alterations in DMN connectivity, including reduced within-DMN coupling post-stress and cortisol-modulated PCC connectivity changes, suggesting an indirect link between HPA axis activity and DMN/ PCC dynamics during aversive experiences. Moreover, acute stress-induced cortisol increases have been shown to correlate with decreased DMN connectivity, including connectivity involving hubs such as the PCC, in large fMRI studies of stress reactivity (Zhang et al., 2019). Taken together, the results on MS C indicate that these modulations are not merely reflecting the task’s cognitive load but may be an important neural mechanism of affective adaptation to adversity in high-resilience individuals.

However, MS C’s dynamic was not the only MS modulation we observed during the VR-ES experience. Consistent with prior findings by Nazare and Tomescu, we observed increased temporal-dynamics parameters of MS D during VR-ES (**Figure 5**), in contrast to neutral resting and positive affective states (Nazare & Tomescu, 2024). As the authors suggested, this result may reflect the negative bias in valence reactivity and the attentional load required for adaptation to aversive experience. As was the case for MS C, we further generalise the MS D findings by showing these mechanisms are independent of the nature of the task, as long as negative valence affective states are reported. These results are also in line with the MSs growing literature showing that MS D dynamics are enhanced during an attentional task, and the neural generators of MS D are localised within the main hubs of the dorsal attention network (DAN) and executive-functional generators involved in cognitive control (Britz et al., 2010; Custo et al., 2017; Tarailis et al., 2024). Thus, our results suggest that prolonged MS D during VR-ES may reflect the functional relevance of the DAN, which is triggered during immersive VR-ES exposure, for focus switching, attentional shifting, and emotion regulation, thereby supporting efficient adaptation and faster recovery. This type of response can be interpreted as an adaptive reactivity strategy, whereby the cognitive system prioritises integrating external stimuli to respond efficiently to perceived threats in VR ES, and it might be reflecting a basic mechanism of adaptation to aversiveness that predicts emotional regulation as a function of psychological resilience traits, and prior aversive experiences, as suggested by our moderation analyses. Indeed, MS D dynamics could interact with individual resilience traits, shaping emotional and anxiety reactivity POST-VR-ES. Our results highlighted that increases in VR-ES MS D were associated with lower emotional reactivity in high-resilience individuals and with reduced anxiety reactivity in low-trauma participants POST-VR-ES, further suggesting that MS D modulation during negative valence affective states might be independent of the endogenous-exogenous nature of the affective stimuli. (**Figure 6bd**).

From an endocrine resilience perspective, we were surprised to find no association of MS D with VR-ES cortisol modulation, as MS D has been previously related to exogenous oxytocin (OXT) (Schiller et al., 2019; Tomescu et al., 2024), and OXT seems to be essential in emotional regulation and stress recovery (Engert et al., 2014). Several studies, including one from our lab using intranasal OXT administration, have found a significant increase in task-positive networks, such as the attention-related MS D. Resting-state activity after oxytocin administration showed a transition from internal to external processing, via decreasing DMN-related MS C and increasing the DAN-associated MS D, with changes varying as a function of inter-individual variability in social functioning and anxiety (Schiller et al., 2019; Tomescu et al., 2024). Taken together, the findings on MS D indicate that its modulation during VR-ES supports information processing in ways that facilitate cognitive control and emotional recovery in individuals with high resilience, highlighting its potential role in adaptive responses.

Finally, we found that MS B decreased during VR-ES reactivity when compared to VR-CTRL. This MS has been associated with regions in the occipital cortex and with autobiographical memory and scene visualisation in task-related studies (Britz et al., 2010; Custo et al., 2017; Tarailis et al., 2024). In affective contexts, increased MS B is mostly observed as decreased during negative states and increased during self-generated positive-valence affective experiences (Nazare & Tomescu, 2024). In our data, consistent with these findings, the occurrence rate of MS B in the VR-ES group is lower in the VR-CTRL group. Given that the VR-CTRL participants experienced a calm, neutral setting, which likely promoted a mildly positive emotional state, this finding may also indicate greater engagement of autobiographical memory and scene visualisation during the VR-CTRL experience than during the VR-ES. Furthermore, resilience and childhood aversive experiences moderate the negative association with emotional recovery. Indeed, reduced MS B microstates were associated with higher affective recovery in highly resilient individuals.

In summary, the findings collectively characterize the neurodynamic and neuroendocrine mechanisms of VR-ES reactivity and its association with emotional regulation as a function of resilience traits and early adversity. In the following paragraph, we will discuss how these large-scale brain dynamics subsequently reorganize during the recovery phase to sustain long-term resilience.

### EEG MS modulations during POST VR-ES resting-state recovery

As distinct neuro-hormonal moments in acute stress might promote long-term resilience (de Kloet & Joëls, 2024), we further investigated resting-state EEG MSs activity during the recovery phase, and our results confirm that distinct neural patterns are activated during this period. In opposition to the VR-ES reactivity phase, during the recovery phase, an increase in MS C and a reduction in MS D were observed, in addition to a reduction in MS E (**Figure 7cd**). The enhancement of MS C during recovery after aversive experience is consistent with its functional role, which reflects top-down brain activity patterns associated with relaxation and stimulus-independent, internally oriented thinking (Faber et al., 2017; Katayama et al., 2007; Milz et al., 2016). In line with this, the observed reduction in microstate D is consistent with its functional role, as this microstate reflects top-down brain activity patterns associated with attentional states and cognitive task engagement (Seitzman et al., 2017). Therefore, we interpret the decrease in its occurrence following VR-ES as a natural disengagement from task-related, externally oriented processing after task completion, and integration of these experiences.

Additionally, our results are partially aligned with the literature. Our findings align with those of Hu et al., who observed increases in temporal descriptors related to MS C (coverage and occurrence rate) following acute stress induced by the TSST (Hu et al., 2021). They proposed that the pronounced increase in MS C after acute stress may reflect stress-induced norepinephrine release, which amplifies salience network activity, suppresses the default mode network, and engages the central executive network. Importantly, Hu et al. reduced the number of microstate maps to four, rather than considering dominant topographies, and under this clustering approach, MS C may partially overlap with salience network activity. However, our MS C, separately from salience-associated MS, also showed increased presence; we confirm their findings, however, the interpretation might point towards more of the experiential integration part of the self-referential activity, associated with DMN activity that supports emotional regulation at different stages of adaptation. Furthermore, as opposed to Hu et al. (2021), we also observed decreases in MSs D and E, suggesting distinct neuronal mechanisms associated with changes in other fMRI networks for natural disaster adverse experiences than during psychosocial stress.

More importantly, Hu et al., 2021 reported significantly larger increases in cortisol compared to our study. This difference is likely attributable to the use of the TSST, a well-established and highly potent social-evaluative stressor known to elicit robust HPA axis activation (Dickens et al., 2026; Liu et al., 2021). In contrast, our immersive VR earthquake experience elicited a comparatively milder physiological cortisol response.

While the overall MS dynamics reveal general recovery patterns, behavioural trait-level factors such as trauma may shape how these neural changes are associated with emotional regulation. Indeed, we found that individuals with low trauma show a positive association between Δ POST-PRE VR-ES MS D mean duration and Δ T_1_-T_0_ emotional reactivity (**Figure 8a**). Thus, the shorter the D duration POST-PRE MS D, the greater the emotional recovery in individuals with low levels of adverse childhood experiences. This finding suggests an adaptive, internally oriented coping process that supports self-referential evaluation and the integration of emotional responses following VR-ES while executive functions are disconnected. Rather than indicating a passive return to a relaxed state, this pattern may index a flexible engagement of MS D DAN & MS C DMN-mediated processing that enables the adaptive integration of exogenous aversive-induced emotions after an earthquake. Indeed, several studies indicate that increased DMN activity may support the integration of external stimuli (Gao et al., 2023; Hasson et al., 2015).

Finally, we found MS E modulations during the POST-VR-ES, which is among the recently investigated microstates and is associated with the salience network (SN). When studies are restricted to four microstates, it may partially overlap with MS C. Custo et al. demonstrated that these two microstates are supported by distinct neural generators, with MS E being linked to activity in the dorsal anterior cingulate cortex, superior frontal gyrus, bilateral middle prefrontal cortex, and insular cortices, serving as hubs of the salience network involved in the processing of interoceptive and emotional information (Custo et al., 2017; Dragu et al., 2025; Tarailis et al., 2024). In our data, MS E showed decreased temporal dynamics, similar to the other task positive network MS D, and in opposition to MS C, increased presence during recovery. Moreover, our results exhibited opposite patterns after VR-ES was significantly negatively correlated, providing additional evidence for their distinct neural generators and functional roles. To our surprise, we did not find any significant modulation of MS E during the VR-ES reactivity, as we would expect based on our previous research, where MS E has been reported to be reduced during negative-valence effective states, and real-life highly arousing scenarios, like a helicopter maneuver or after iTBS stimulations promoting lDLPFC activity (Deolindo et al., 2021; Dragu et al., 2025; Nazare & Tomescu, 2024; Tarailis et al., 2024). This might suggest that SN-related MS E modulation during self-generated negative affective states and during recovery after VR-ES might be more involved in a shift from externally oriented salience detection toward the internal evaluation of affective states and the interoceptive process of integrating the aversive experience into a meaningful experience and self-referential evaluation and integration of emotional responses.

### Association between salivary cortisol changes and MS dynamics

Salivary cortisol, a hormone known to play an essential role in the acute stress response and long-term resilience, has been shown to increase following exposure to acute psychosocial stressors such as the TSST (de Kloet & Joëls, 2024; Narvaez Linares et al., 2020). In our study, we showed a relatively small elevation in salivary cortisol levels in the VR-ES group compared with the VR-CTRL group. We typically observe a much greater increase in cortisol levels following such psychosocial stress tasks as the TSST. However, important limitations of the TSST include its ecological validity and labour-intensive nature; it is not the scope of this paper to compare these results. Moreover, in our study, for ethical reasons, participants were informed in both conditions that they would undergo an aversive VR experience and that they were free to leave at any time. No participant aborted the VR-ES; however, the high level of cortisol in the beginning might be explained by their anticipation of aversiveness. Nevertheless, to better understand neurohormonal dynamics, we examined results separately for cortisol responders and non-responders during the reactivity phase. Cortisol responders had a significantly lower incidence than nonresponders in MS C during VR-ES (**Figure 9a**), thus confirming that the decrease in MS C during VR-ES is part of an adaptive neuro-hormonal mechanism. This suggests a possible coupling between DMN-related MS C and cortisol release that sustains emotional regulation, further indicating that MS C modulations during reactivity are not merely reflecting the task’s cognitive load but may reflect an important neural mechanism of affective adaptation to natural disaster adversity in high-resilience individuals.

Importantly, these neurohormonal mechanisms extend beyond VR-ES reactivity and contribute to the reorganization of microstate dynamics during the recovery phase. In line with Kloet & Joëls , our results also support the idea that cortisol is a marker of resilience rather than vulnerability, with higher cortisol levels and neural activity predicting better emotional recovery and long-term resilience.

While these findings suggest a potential modulatory role of cortisol in aversive experiences-related microstate dynamics, their possible causal effect should be interpreted cautiously, as larger samples are needed to substantiate and elucidate the role of salivary cortisol in shaping microstate activity patterns that promote resilience to natural disasters.

### Conclusions and future perspectives

By investigating EEG modulations following an immersive VR-simulated earthquake exposure, we demonstrated that VR can be used to study adaptation to natural disasters. Moreover, we identified an adaptive neurohormonal mechanism in response to extreme environmental threats, moderated by general resilience traits and early-life adverse experiences. Specifically, our results indicate that MSs C, D, and E index distinct phases of neural reactivity and recovery in individuals with high resilience and low trauma experience. Most importantly, these results indicate that accurate inferences about the mechanisms underlying reactivity to aversive experiences can be drawn only by investigating the aftereffects of aversive events, as our results show that these effects can yield completely reversed patterns of activity. Characterizing such adaptive neural signatures may constitute a central framework for subsequent investigations under VR extreme environmental conditions.

However, several limitations of the present study should be noted. Although the study includes both an experimental and a control group, the small sample size is a limitation, and the focus on resilience should also extend to lower levels of resilience beyond the academic student population. Another limitation concerns the lack of assessment of presence and immersion during the VR exposure, as no standardized questionnaires were administered to quantify these subjective states. Future studies should integrate these measures to better elucidate the relationship between the temporal dynamics of microstates and the cognitive processes that may be modulated by VR exposure simulating a natural disaster.

Although this was a first step toward studying mobile brain EEG adaptation to environmental challenges, we need to examine other types of natural disasters and use finer-grained temporal resolution, such as real-time analysis, to further increase the generalizability of these findings. However, we believe these results are already highly important for understanding the neurohormonal mechanisms underlying acute adaptation to environmental challenges and for informing targeted interventions to sustain resilience in highly affected populations.

## Acknowledgements

We thank Sabin Șerban and Dan Făcăeru for their essential role in the development of the EEG mobile system, implementation and synchronization of the EEG-VR CAVE systems. We would also like to acknowledge the UNATC University, LDCAPEI lab for fostering the NEURESIL project, the lab director Prof. Alexandru Berceanu for his constant support, the administration and university rector Prof. Liviu Lucaci.

## REFERENCES

Akounach, M., Yigitalp, B. M., Kizilisik, S., Duman, D., Aarabi, A., Lelard, T., & Mouras, H. (2026). Neural signatures of environmental perception: EEG and postural markers differentiate responses to polluted vs. pleasant landscapes. Journal of Environmental Psychology, 111, 103001. 10.1016/j.jenvp.2026.103001

Antonova, E., Holding, M., Suen, H. C., Sumich, A., Maex, R., & Nehaniv, C. (2022). EEG microstates: Functional significance and short-term test-retest reliability. Neuroimage: Reports, 2(2). 10.1016/j.ynirp.2022.100089

Artoni, F., & Michel, C. M. (2025). How does Independent Component Analysis Preprocessing Affect EEG Microstates? Brain Topography, 38(2). 10.1007/s10548-024-01098-4

Ascone, L., Mostajeran, F., Mascherek, A., Tawil, N., Knaust, T., Samaan, L., & Kühn, S. (2025). Multi-vs. unimodal forest-bathing in VR to enhance affective and cognitive recovery after acute stress. Journal of Environmental Psychology, 105. 10.1016/j.jenvp.2025.102637

Banholzer, S., Kossin, J., & Donner, S. (2014). The impact of climate change on natural disasters. In Reducing Disaster: Early Warning Systems for Climate Change (Vol. 9789401785983, pp. 21–49). Springer Netherlands. 10.1007/978-94-017-8598-3_2

Benjamini, Y. (1995). Controlling The False Discovery Rate-A Practical And Powerful Approach To Multiple Testing. Article in Journal of the Royal Statistical Society Series B. 10.2307/2346101

Bernstein, D. P., Stein, J. A., Newcomb, M. D., Walker, E., Pogge, D., Ahluvalia, T., Stokes, J., Handelsman, L., Medrano, M., Desmond, D., & Zule, W. (2003). Development and validation of a brief screening version of the Childhood Trauma Questionnaire. Child Abuse and Neglect, 27(2), 169–190. 10.1016/S0145-2134(02)00541-0

Britz, J., Van De Ville, D., & Michel, C. M. (2010). BOLD correlates of EEG topography reveal rapid resting-state network dynamics. In NeuroImage (Vol. 52, Number 4, pp. 1162–1170). Academic Press Inc. 10.1016/j.neuroimage.2010.02.052

Brunet, D., Murray, M. M., & Michel, C. M. (2011). Spatiotemporal analysis of multichannel EEG: CARTOOL. In Computational Intelligence and Neuroscience (Vol. 2011). 10.1155/2011/813870

Chivu, A., Pascal, S. A., Damborská, A., & Tomescu, M. I. (2024). EEG Microstates in Mood and Anxiety Disorders: A Meta-analysis. Brain Topography, 37(3), 357–368. 10.1007/s10548-023-00999-0

Cohen, J. (1988). Statistical Power Analysis for the Behavioral Sciences Second Edition.

Connor, K. M., & Davidson, J. R. T. (2003). Development of a new Resilience scale: The Connor-Davidson Resilience scale (CD-RISC). Depression and Anxiety, 18(2), 76–82. 10.1002/da.10113

Custo, A., Van De Ville, D., Wells, W. M., Tomescu, M. I., Brunet, D., & Michel, C. M. (2017). Electroencephalographic Resting-State Networks: Source Localization of Microstates. Brain Connectivity, 7(10), 671–682. 10.1089/brain.2016.0476

Daube, A., Lima-Carmona, Y. E., Hernández Solís, D. G., & Contreras-Vidal, J. L. (2026). A Systematic Review and Meta-Analysis of EEG, fMRI, and fNIRS Studies on the Psychological Impact of Nature on Well-Being. International Journal of Environmental Research and Public Health, 23(3), 377. 10.3390/ijerph23030377

de Kloet, E. R., & Joëls, M. (2024). The cortisol switch between vulnerability and resilience. In Molecular Psychiatry (Vol. 29, Number 1, pp. 20–34). Springer Nature. 10.1038/s41380-022-01934-8

Deng, K., Xing, S., Wang, G., Hu, X., & Chen, T. (2024). A Clarity-intensity model for evacuation panic by fNIRS and VR. Journal of Environmental Psychology, 93. 10.1016/j.jenvp.2023.102228

Denzer, S., Diezig, S., Achermann, P., Mast, F. W., & Koenig, T. (2024). Electrophysiological (EEG) microstates during dream-like bizarre experiences in a naturalistic scenario using immersive virtual reality. European Journal of Neuroscience, 60(8), 5815–5830. 10.1111/ejn.16530

Deolindo, C. S., Ribeiro, M. W., de Aratanha, M. A. A., Scarpari, J. R. S., Forster, C. H. Q., da Silva, R. G. A., Machado, B. S., Amaro Junior, E., König, T., & Kozasa, E. H. (2021). Microstates in complex and dynamical environments: Unraveling situational awareness in critical helicopter landing maneuvers. Human Brain Mapping, 42(10), 3168–3181. 10.1002/hbm.25426

Dickens, H., Gournay-Berman, L. R., Hill, M., Joslin, M. D. M., Walker, J., Makhanova, A., Leen-Feldner, E., & Vargas, I. (2026). Hypothalamic-pituitary-adrenal (HPA) axis stress response to a laboratory-based 10 % CO2 challenge in healthy adults. Psychoneuroendocrinology, 184. 10.1016/j.psyneuen.2025.107712

Dowdy, A. J., Ye, H., Pepler, A., Thatcher, M., Osbrough, S. L., Evans, J. P., Di Virgilio, G., & McCarthy, N. (2019). Future changes in extreme weather and pyroconvection risk factors for Australian wildfires. Scientific Reports, 9(1). 10.1038/s41598-019-46362-x

Dragu, M. A., Niculescu, G., & Tomescu, M. I. (2025). EEG Microstates Signatures of rTMS Response Over the lDLPFC: A Band-Specific Analysis. Brain Topography, 38(6). 10.1007/s10548-025-01146-7

Engert, V., Smallwood, J., & Singer, T. (2014). Mind your thoughts: Associations between self-generated thoughts and stress-induced and baseline levels of cortisol and alpha-amylase. Biological Psychology, 103, 283–291. 10.1016/j.biopsycho.2014.10.004

Faber, P. L., Travis, F., Milz, P., & Parim, N. (2017). EEG microstates during different phases of Transcendental Meditation practice. Cognitive Processing, 18(3), 307–314. 10.1007/s10339-017-0812-y

Fritz, C. O., Morris, P. E., & Richler, J. J. (2012). Effect size estimates: Current use, calculations, and interpretation. Journal of Experimental Psychology: General, 141(1), 2–18. 10.1037/a0024338

Gao, H., Zeng, Y., Tian, T., Liu, C., Wu, J., Wu, H., & Qin, S. (2023). Long-term stress shapes dynamic reconfiguration of functional brain networks across multi-task demands. 10.1101/2023.03.28.534193

Gramann, K. (2024). Mobile EEG for neurourbanism research - What could possibly go wrong? A critical review with guidelines. In Journal of Environmental Psychology (Vol. 96). Academic Press. 10.1016/j.jenvp.2024.102308

Gramfort, A., Luessi, M., Larson, E., Engemann, D. A., Strohmeier, D., Brodbeck, C., Goj, R., Jas, M., Brooks, T., Parkkonen, L., & Hämäläinen, M. (2013). MEG and EEG data analysis with MNE-Python. Frontiers in Neuroscience, (7 DEC). 10.3389/fnins.2013.00267

Hameed, A., Möller, S., & Perkis, A. (2023). How good are virtual hands? Influences of input modality on motor tasks in virtual reality. Journal of Environmental Psychology, 92. 10.1016/j.jenvp.2023.102137

Hanshans, C., Amler, T., Zauner, J., & Bröll, L. (2024). Inducing and measuring acute stress in virtual reality: Evaluation of canonical physiological stress markers and measuring methods. Journal of Environmental Psychology, 94. 10.1016/j.jenvp.2023.102107

Hasson, U., Chen, J., & Honey, C. J. (2015). Hierarchical process memory: Memory as an integral component of information processing. In Trends in Cognitive Sciences (Vol. 19, Number 6, pp. 304–313). Elsevier Ltd. 10.1016/j.tics.2015.04.006

Hu, N., Long, Q., Li, Q., Hu, X., Li, Y., Zhang, S., Chen, A., Huo, R., Liu, J., & Wang, X. (2021). The modulation of salience and central executive networks by acute stress in healthy males: An EEG microstates study. International Journal of Psychophysiology, 169, 63–70. 10.1016/j.ijpsycho.2021.09.001

Huijsmans, M. K., Rieder, L., Kreuer, K., & Müller, B. C. N. (2025). Less is more - The effect of VR-induced awe on minimalism and sustainable consumption. Journal of Environmental Psychology, 105. 10.1016/j.jenvp.2025.102685

Katayama, H., Gianotti, L. R. R., Isotani, T., Faber, P. L., Sasada, K., Kinoshita, T., & Lehmann, D. (2007). Classes of multichannel EEG microstates in light and deep hypnotic conditions. Brain Topography, 20(1), 7–14. 10.1007/s10548-007-0024-3

Kawai, C., Georgiou, F., Pieren, R., Tobias, S., Mavros, P., & Schäffer, B. (2024). Investigating effect chains from cognitive and noise-induced short-term stress build-up to restoration in an urban or nature setting using 360° VR. Journal of Environmental Psychology, 100. 10.1016/j.jenvp.2024.102466

Kerby, D. S. (2014). The Simple Difference Formula: An Approach to Teaching Nonparametric Correlation. Comprehensive Psychology, 3, 11.IT.3.1. 10.2466/11.it.3.1

Kim, K., Duc, N. T., Choi, M., & Lee, B. (2021). EEG microstate features according to performance on a mental arithmetic task. Scientific Reports, 11(1). 10.1038/s41598-020-79423-7

Klug, M., Berg, T., & Gramann, K. (2024). Optimizing EEG ICA decomposition with data cleaning in stationary and mobile experiments. Scientific Reports, 14(1). 10.1038/s41598-024-64919-3

Knez, I., Willander, J., Butler, A., Sang, Å. O., Sarlöv-Herlin, I., & Åkerskog, A. (2021). I can still see, hear and smell the fire: Cognitive, emotional and personal consequences of a natural disaster, and the impact of evacuation. Journal of Environmental Psychology, 74. 10.1016/j.jenvp.2021.101554

Lehmann, D., & Skrandies, W. (1980). Reference-free identification of components of checkerboard-evoked multichannel potential fields. In Electroencephalography and Clinical Neurophysiology (Vol. 48).

Li, H., Shin, H., Sentis, L., Siu, K. C., Millán, J. del R., & Lu, N. (2024). Combining VR with electroencephalography as a frontier of brain-computer interfaces. In Device (Vol. 2, Number 6). Cell Press. 10.1016/j.device.2024.100425

Liu, Q., Wu, J., Zhang, L., Sun, X., Guan, Q., & Yao, Z. (2021). The Relationship Between Perceived Control and Hypothalamic–Pituitary–Adrenal Axis Reactivity to the Trier Social Stress Test in Healthy Young Adults. Frontiers in Psychology, 12. 10.3389/fpsyg.2021.683914

Ma, H., Zhao, J., Hommel, B., & Ma, K. (2025). Virtual Reality Experience as Reflected in EEG Microstates. Brain Topography, 38(6). 10.1007/s10548-025-01155-6

McGrath, R. E., & Meyer, G. J. (2006). When effect sizes disagree: The case of r and d. Psychological Methods, 11(4), 386–401. 10.1037/1082-989X.11.4.386

Michel, C. M., & Koenig, T. (2018). EEG microstates as a tool for studying the temporal dynamics of whole-brain neuronal networks: A review. In NeuroImage (Vol. 180, pp. 577–593). Academic Press Inc. 10.1016/j.neuroimage.2017.11.062

Miller, R., & Plessow, F. (2013). Transformation techniques for cross-sectional and longitudinal endocrine data: Application to salivary cortisol concentrations. Psychoneuroendocrinology, 38(6), 941–946. 10.1016/j.psyneuen.2012.09.013

Milz, P., Faber, P. L., Lehmann, D., Koenig, T., Kochi, K., & Pascual-Marqui, R. D. (2016). The functional significance of EEG microstates-Associations with modalities of thinking. NeuroImage, 125, 643–656. 10.1016/j.neuroimage.2015.08.023

Mioni, G., & Pazzaglia, F. (2023). Time perception in naturalistic and urban immersive virtual reality environments. Journal of Environmental Psychology, 90. 10.1016/j.jenvp.2023.102105

Mishra, A., Englitz, B., & Cohen, M. X. (2020). EEG microstates as a continuous phenomenon. NeuroImage, 208. 10.1016/j.neuroimage.2019.116454

Mortimer, A., Davies, K., Smith, G., & Ahmed, I. (2026). How lived experiences of disaster displacement reshape place attachment. Journal of Environmental Psychology, 102915. 10.1016/j.jenvp.2026.102915

Murray, M. M., Brunet, D., & Michel, C. M. (2008). Topographic ERP analyses: A step-by-step tutorial review. In Brain Topography (Vol. 20, Number 4, pp. 249–264). 10.1007/s10548-008-0054-5

Musso, F., Brinkmeyer, J., Mobascher, A., Warbrick, T., & Winterer, G. (2010). Spontaneous brain activity and EEG microstates. A novel EEG/fMRI analysis approach to explore resting-state networks. In NeuroImage (Vol. 52, Number 4, pp. 1149–1161). Academic Press Inc. 10.1016/j.neuroimage.2010.01.093

Nagabhushan Kalburgi, S., Kleinert, T., Aryan, D., Nash, K., Schiller, B., & Koenig, T. (2024). MICROSTATELAB: The EEGLAB Toolbox for Resting-State Microstate Analysis. Brain Topography, 37(4), 621–645. 10.1007/s10548-023-01003-5

Narvaez Linares, N. F., Charron, V., Ouimet, A. J., Labelle, P. R., & Plamondon, H. (2020). A systematic review of the Trier Social Stress Test methodology: Issues in promoting study comparison and replicable research. In Neurobiology of Stress (Vol. 13). Elsevier Inc. 10.1016/j.ynstr.2020.100235

Nazare, K., & Tomescu, M. I. (2024). Valence-specific EEG microstate modulations during self-generated affective states. Frontiers in Psychology, 15. 10.3389/fpsyg.2024.1300416

Nuojua, S., Pahl, S., Wyles, K. J., & Thompson, R. C. (2024). The impact of virtual reality exposure on ocean connectedness and consumer responses to single-use packaging. Journal of Environmental Psychology, 99. 10.1016/j.jenvp.2024.102450

Oleksy, T., Lassota, I., Wnuk, A., & Wcześniak, R. (2024). Virtual changes in real places: Understanding the role of place attachment in augmented reality adoption. Journal of Environmental Psychology, 98. 10.1016/j.jenvp.2024.102386

Parsons, T. D. (2015). Virtual reality for enhanced ecological validity and experimental control in the clinical, affective and social neurosciences. Frontiers in Human Neuroscience, 9(DEC). 10.3389/fnhum.2015.00660

Pion-Tonachini, L., Kreutz-Delgado, K., & Makeig, S. (2019). ICLabel: An automated electroencephalographic independent component classifier, dataset, and website. NeuroImage, 198, 181–197. 10.1016/j.neuroimage.2019.05.026

Poulsen, A. T., Pedroni, A., Langer, N., & Hansen, L. K. (2018). Microstate EEGlab toolbox: An introductory guide. 10.1101/289850

Satpute, A. B., & Lindquist, K. A. (2019). The Default Mode Network’s Role in Discrete Emotion. In Trends in Cognitive Sciences (Vol. 23, Number 10, pp. 851–864). Elsevier Ltd. 10.1016/j.tics.2019.07.003

Schiller, B., Koenig, T., & Heinrichs, M. (2019). Oxytocin modulates the temporal dynamics of resting EEG networks. Scientific Reports, 9(1). 10.1038/s41598-019-49636-6

Schiller, B., Sperl, M. F. J., Kleinert, T., Nash, K., & Gianotti, L. R. R. (2024). EEG Microstates in Social and Affective Neuroscience. In Brain Topography (Vol. 37, Number 4, pp. 479–495). Springer. 10.1007/s10548-023-00987-4

Schöne, B., Kisker, J., Lange, L., Gruber, T., Sylvester, S., & Osinsky, R. (2023). The reality of virtual reality. Frontiers in Psychology, 14. 10.3389/fpsyg.2023.1093014

Seitzman, B. A., Abell, M., Bartley, S. C., Erickson, M. A., Bolbecker, A. R., & Hetrick, W. P. (2017). Cognitive manipulation of brain electric microstates. NeuroImage, 146, 533–543. 10.1016/j.neuroimage.2016.10.002

Tarailis, P., Koenig, T., Michel, C. M., & Griškova-Bulanova, I. (2024). The Functional Aspects of Resting EEG Microstates: A Systematic Review. In Brain Topography (Vol. 37, Number 2, pp. 181–217). Springer. 10.1007/s10548-023-00958-9

Tomescu, M. I., Papasteri, C. C., Sofonea, A., Boldasu, R., Kebets, V., Pistol, C. A. D., Poalelungi, C., Benescu, V., Podina, I. R., Nedelcea, C. I., Berceanu, A. I., & Carcea, I. (2022). Spontaneous thought and microstate activity modulation by social imitation. NeuroImage, 249. 10.1016/j.neuroimage.2022.118878

Tomescu, M. I., Van der Donck, S., Perisanu, E. M., Berceanu, A. I., Alaerts, K., Boets, B., & Carcea, I. (2024). Social functioning predicts individual changes in EEG microstates following intranasal oxytocin administration: A double-blind, cross-over randomized clinical trial. Psychophysiology, 61(8). 10.1111/psyp.14581

Vardy, T., & Atkinson, Q. D. (2019). Property Damage and Exposure to Other People in Distress Differentially Predict Prosocial Behavior After a Natural Disaster. Psychological Science, 30(4), 563–575. 10.1177/0956797619826972

Waters, L., Algoe, S. B., Dutton, J., Emmons, R., Fredrickson, B. L., Heaphy, E., Moskowitz, J. T., Neff, K., Niemiec, R., Pury, C., & Steger, M. (2022). Positive psychology in a pandemic: buffering, bolstering, and building mental health. Journal of Positive Psychology, 17(3), 303–323. 10.1080/17439760.2021.1871945

Wallemacq, P., Below, R., & McClean, D. (2018). Economic losses, poverty & disasters: 1998-2017. United nations Office for disaster risk Reduction.

Yuan, H., Zotev, V., Phillips, R., Drevets, W. C., & Bodurka, J. (2012). Spatiotemporal dynamics of the brain at rest - Exploring EEG microstates as electrophysiological signatures of BOLD resting state networks. NeuroImage, 60(4), 2062–2072. 10.1016/j.neuroimage.2012.02.031

Zanier, E. R., Zoerle, T., Di Lernia, D., & Riva, G. (2018). Virtual reality for traumatic brain injury. Frontiers in Neurology, 9(MAY). 10.3389/fneur.2018.00345

Zhang, W., Hashemi, M. M., Kaldewaij, R., Koch, S. B. J., Beckmann, C., Klumpers, F., & Roelofs, K. (2019). Acute stress alters the ‘default’ brain processing. NeuroImage, 189, 870–877. 10.1016/j.neuroimage.2019.01.063

Zhao, Q., Prooijen, J. W. van, & Spadaro, G. (2024). Coping capacity attenuates the effect of natural disaster risk on conspiracy beliefs. Journal of Environmental Psychology, 97. 10.1016/j.jenvp.2024.102363

